# Contingency and selection in mitochondrial genome dynamics

**DOI:** 10.1101/2021.11.15.468706

**Authors:** Christopher J. Nunn, Sidhartha Goyal

## Abstract

Eukaryotic cells contain numerous copies of mitochondrial DNA (mtDNA), allowing for the coexistence of mutant and wild-type mtDNA in individual cells. The fate of mutant mtDNA depends on their relative replicative fitness within cells and the resulting cellular fitness within populations of cells. Yet the dynamics of the generation of mutant mtDNA and features that inform their fitness remain unaddressed. Here we utilize long read single-molecule sequencing to track mtDNA mutational trajectories in *Saccharomyces cerevisiae*. We show a previously unseen pattern that constrains subsequent excision events in mtDNA fragmentation. We also provide evidence for the generation of rare and contentious non-periodic mtDNA structures that lead to persistent diversity within individual cells. Finally, we show that measurements of relative fitness of mtDNA fit a phenomenological model that highlights important biophysical parameters governing mtDNA fitness. Altogether, our study provides techniques and insights into the dynamics of large structural changes in genomes that may be applicable in more complex organisms.

## Introduction

The mitochondrial genome (mtDNA) in eukaryotic cells encodes a subset of enzymes involved in cellular respiration. Interestingly, the integrity of mtDNA has been implicated in critical biological processes other than respiration such as in apoptosis, trace element and intermediary metabolism, heme synthesis, and iron-sulfur cluster biogenesis (Veatch et al., 2009). Because mtDNA exists in multiple copies within numerous mitochondrial compartments, localized mtDNA damage produces heteroplasmic states with coexisting mutant and wild-type mtDNA in cells. Both intracellular mtDNA dynamics and intercellular selection then ultimately shape the fate of cell populations, with mtDNA damage resulting in cellular defects in single-celled organisms such a yeast and aging and disease in multicellular organisms such as humans (Fayet et al., 2002; Chan et al., 2006; Payne & Chinnery, 2015).

In yeast, which is an excellent model system to study mtDNA dynamics due to the dispensability of mtDNA, mtDNA integrity was linked to the Petite phenotype (Ephrussi et al., 1949). The Petite phenotype is due to destructive recombination events in mtDNA that excise portions of the wild-type genome (Bernardi et al., 1975). These sub-genomic mtDNA fragments are amplified in Petite cells, leaving Petites with the inability to respire due to the loss of genes involved in cellular respiration that exist in the intact wild-type genome. Much has been known about the structure of mtDNA in Petite cells, such as the basis of the excision mechanism that generates their characteristic incomplete mtDNAs and the source of the replication advantage over wild-type mtDNA that is responsible for their existence. But important questions about mtDNA dynamics remain unanswered. These include understanding the rules of the generation of new Petite mtDNA from existing Petite mtDNA, as well as explaining the structure-function relationship in the out-competition of wild-type mtDNA by Petite mtDNA.

Early studies of Petite mtDNA structure revealed that segments excised from the wild-type genome existed as tandem repeats in concatemer structures of various lengths (Locker et al., 1974; Locker et al., 1979; Faugeron-Fonty et al., 1979), with a few debated observations of nonperiodic structures (Heyting et al., 1979; Bos et al., 1980; Faugeron-Fonty et al., 1983). This led to a remarkable collection of work in structural genomics that detailed the excision mechanism - sequence-specific illegitimate recombination within and between abundant repeated AT regions and GC clusters in non-coding regions (Bernardi et al., 1976; Bernardi & Bernardi, 1980; Marotta et al., 1982; de Zamaroczy et al., 1983). It was also revealed that some Petite strains exhibited heterogeneity in mtDNA structure (Bernardi et al., 1976; Lewin et al., 1978; Lewin et al., 1979; Locker et al., 1979), pointing to a dynamic process with continued mtDNA excision events in Petites. This was followed by a collection of studies with the goals of understanding the potential roles of the abundant non-coding sequences involved in excisions and the locations and structure of the mitochondrial origins of replication. These studies showed that most spontaneous Petite mtDNAs were derived from a diverse set of locations in the wild-type genome. It was also shown that their sub-genomic mtDNA fragments contained either one of eight origins of replication (de Zamaroczy et al., 1979; de Zamaroczy et al., 1981), or surrogate replication origins (Goursot et al., 1982).

Given the exclusive presence of replication origins in Petite mtDNA and the observation that Petite mtDNAs with exceedingly small genomes consistently outcompeted wild-type genomes, it was suggested that the replication advantage of Petite mtDNAs was afforded by a higher density of replication origins (Goursot et al., 1980; Blanc & Dujon., 1980; de Zamaroczy et al., 1981). Studies proceeded to investigate the relationship between mtDNA structure and suppressivity, which is a measure of the replication advantage of Petite mtDNA over Grande (wild-type) mtDNA in a cross and was critical to understanding the propagation of Petite mtDNAs. After analyzing a large sample of spontaneous Petites mtDNAs, it was shown that in most cases suppressivity was correlated with origin density and that suppressivity was reduced when canonical replication origins were disrupted or absent (de Zamaroczy et al., 1981; Mangin et al., 1983). Notable exceptions to these rules were described in (Rayko et al., 1988), revealing an even more complex structure-suppressivity relationship.

Near the time of the complete sequencing of the mitochondrial genome of yeast in 1998 (Foury et al., 1998), work on the minute structural details and functional questions regarding mtDNA in Petites appeared to wane. However, a number of questions about the dynamics of Petite mtDNAs remained to be explored fully: (1) understanding the empirical distribution of the locations of excision events that generate Petite genomes, which are influenced both by varying generative frequency at different locations and selection for origin containing fragments, (2) detecting and exploring excision cascades (continued excisions) in existing Petites strains, (3) understanding the generation of rare non-periodic mtDNA structures, (4) resolving heteroplasmy vs homoplasmy in producing the persistent observed structural heterogeneity in Petite strains, and (5) understanding what components of a genome replication model are necessary to capture how mtDNA structure informs suppressivity.

In this study, we address each of these questions which are motivated by an old problem, but with new long read sequencing technology and accompanying structural inference methods. We highlight some advantages of Nanopore sequencing in addressing these questions and future ones, but also technical challenges specific to structure reconstruction with Nanopore sequencing of the mtDNA in yeast. Among contributions to all of the aforementioned questions, we showcase a previously unseen pattern that constrains subsequent excision events in generating new Petite mtDNA structures from existing ones, settle contention in the literature surrounding the existence and generation of non-periodic “mixed” Petite structures, and propose a phenomenological model of suppressivity that highlights important biophysical parameters governing mtDNA fitness.

## Results

### Characterizing excision events that results in petites

#### Overview of the structure of Grande and Petite mtDNA

To quantify mtDNA structure and their dynamics we opted to sequence both Petite and Grande colonies with a Nanopore MinION single molecule sequencing platform. We expected the long reads generated from this sequencing technology to improve structure reconstruction for both high and low frequency structures compared to short read sequencing approaches. In total we sequenced 38 Petite colonies derived from 9 spontaneous Petite colonies through passaging and 10 Grande (wild-type) colonies of the same *S. cerevisiae* strain. Starting with 9 spontaneous Petite colonies, each colony was passaged twice onto new media, storing and culturing three colonies at each passage. This generated families of Petite colonies, with 9 colonies sharing each spontaneous progenitor after two passages (Figure 1a). The suppressivities of all colonies sharing a progenitor were measured (see methods), but only a subset were sequenced (dotted circles in Figure 1a). Subfamilies, labeled in Figure 1a as a subscript, were grouped based on differing mtDNA content from other members in the same family. The coverage curves from the sequencing of each colony are shown in Figure 1b and provide a coarse picture of their mtDNA content. It is evident that some colonies within families, such as family 4, have differing mtDNA content but share the same spontaneous Petite colony progenitor. This is due to either ongoing mtDNA changes during growth, or mtDNA diversity within spontaneous progenitor colonies that segregate into different cells through genome bottlenecks during budding. However, comparing the mtDNA content in colonies that share second passage progenitors reveals that two passages followed by culturing (~32 generations) was sufficient to homogenize mtDNA content in all cases except family 1.

**Figure 1:**
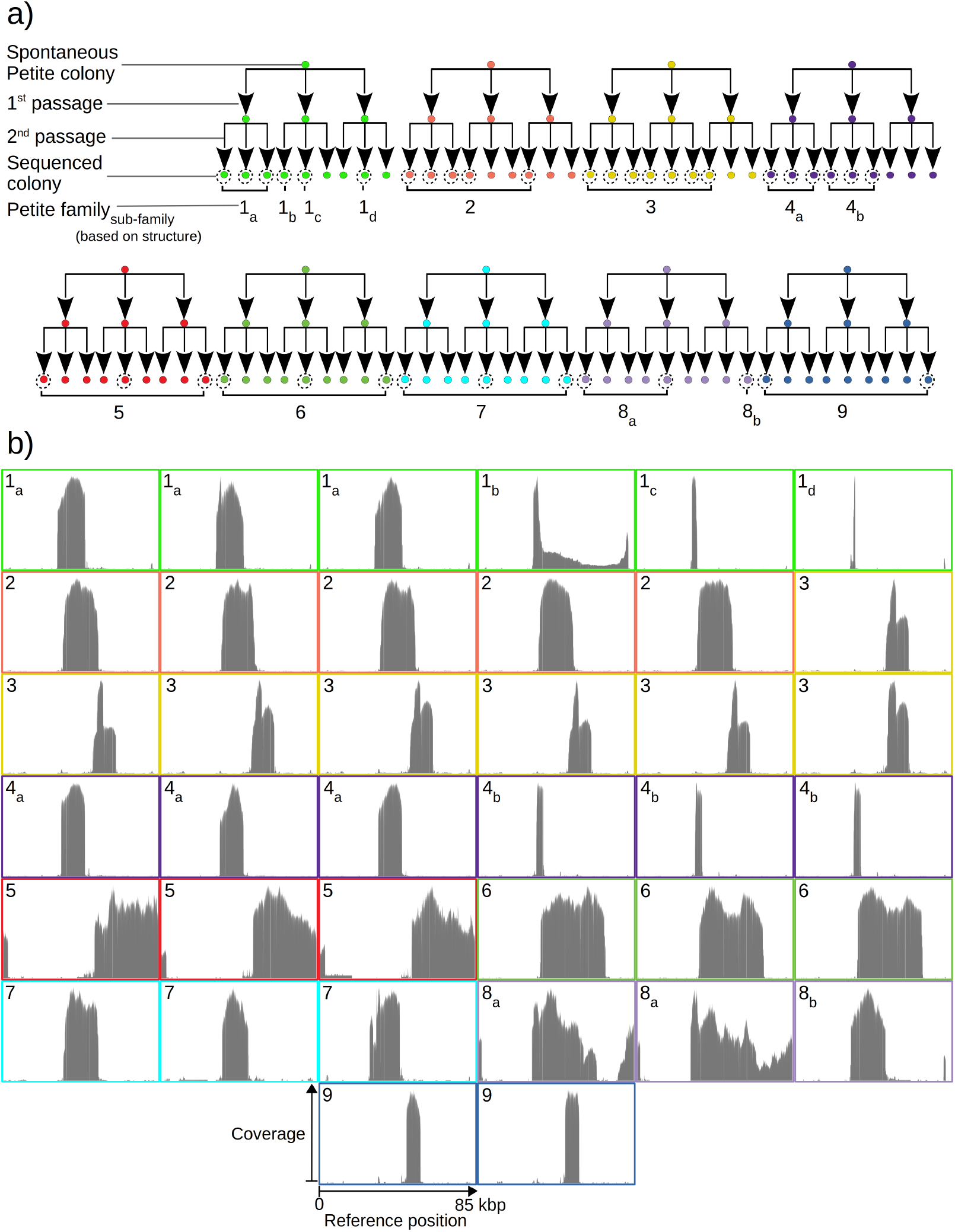
a) An overview of the architecture of the Petite colony sequencing experiment in this study. Nine spontaneous Petite colonies were passaged twice onto new media, culturing and storing three colonies for each passage. This produced families of colonies (indicated by colour), where all colonies after two passages were derived from the same spontaneous Petite colony progenitor, but only a subset of colonies were cultured and then sequenced with a Nanopore MinION sequencing device (indicated by dotted circles). In addition to families, sub-families are labeled as a subscript and grouped based on the predominant mtDNA structure present in these colonies according to sequencing results. The sequencing coverage (arbitrary coverage scaling, consistent reference location scaling) is shown in panel (b).

Mapping of the mtDNA to a reference sequence, followed by careful filtering of inverted duplication artifacts (Figure S1) and clustering of alignment breakpoint signals with a variety of parameters (see methods), revealed both inverted and non-inverted mtDNA breakpoints in all Petite colonies and rare mtDNA breakpoints in Grande colonies. These breakpoint signals delineate sequence alignments that are collinear with the reference mtDNA sequence, but merged in such a way that disjoint alignment locations on the reference genome have been brought together. Non-inverted breakpoints indicate the merging of disjoint sequences in the reference from the same strand, or with the same orientation, while inverted breakpoints indicate the merging of disjoint mtDNA sequences on opposite strands (Figure 2a). Long reads with an average read length of 6kbp and maximum length of 120kbp directly revealed that these breakpoint signals were contributed by concatemer structures in Petites, composed of tandem repeats of sub-genome sized repeat units that had been excised from the wild-type genome and amplified into repeated structures (Figure 2b). Grande reads also revealed concatemer structures manifested in reads as subsamples of genome sized repeating units devoid of breakpoint signals (Figure 2c). These concatemer structures in Petites with sub-genome sized repeat units are consistent with the existing literature on mtDNA structure in Petites that relied on restriction digestion mapping and electron microscopy (Bernardi et al., 1976; Lewin et al., 1978; Locker et al., 1979; Faugeron-Fonty et al., 1979; Bernardi & Bernardi, 1980).

**Figure 2:**
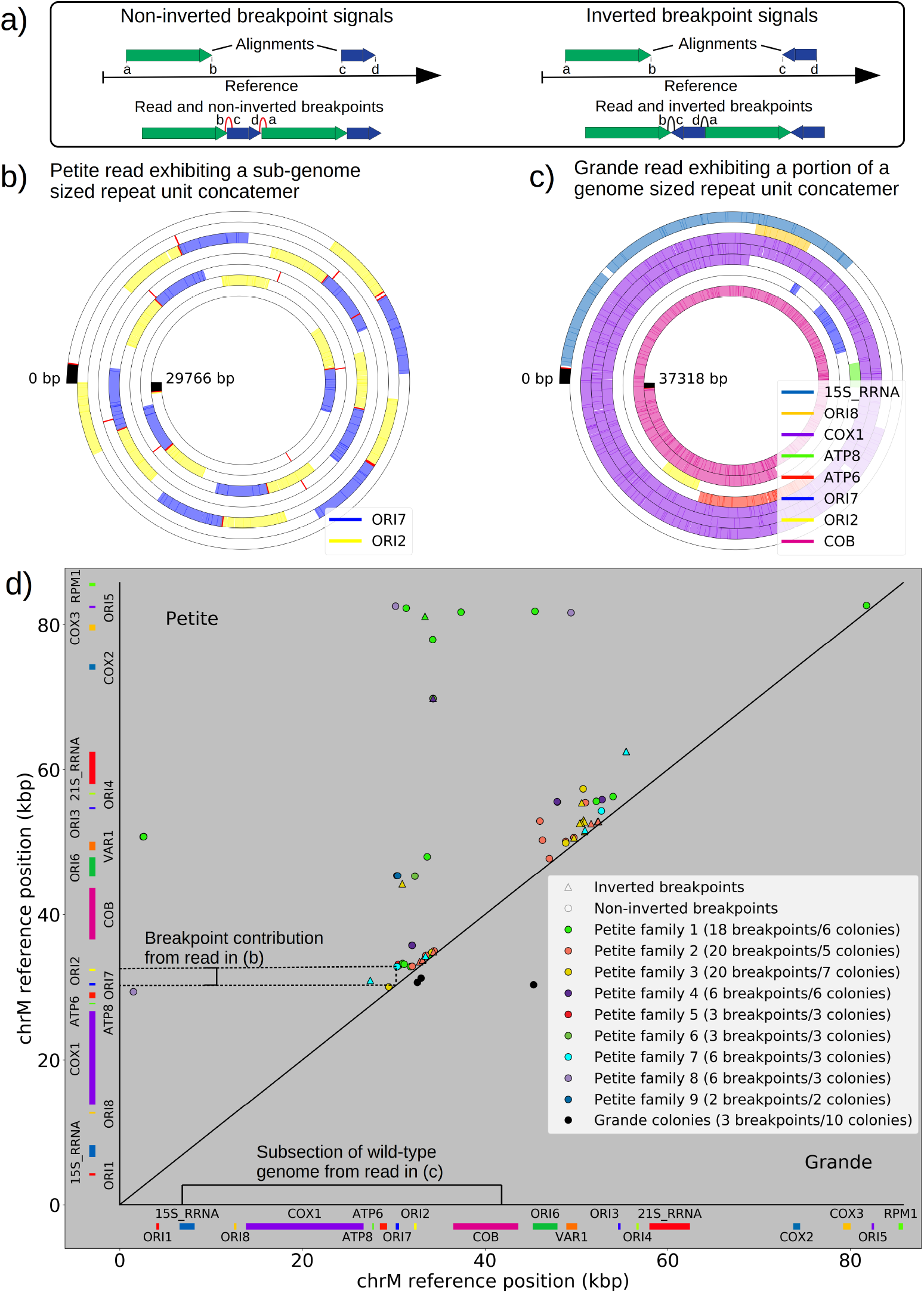
a) A schematic of the definition of alignments and breakpoint signals. Alignments (which are sequences collinear with the reference genome) with their location on the reference are shown as coloured arrows alongside the coordinates of alignment edges (a,b) and (c,d). A hypothetical read is shown below these alignments, indicating how the alignments are oriented with respect to each other and the coordinates of the alignment edges in contact that define the breakpoints denoted as arcs. Non-inverted breakpoints represent merged alignments from disjoint locations on the reference that map to the same strand of DNA (red arcs), while inverted breakpoints represent the same disjoint merging of alignments but with different orientation (black arcs). b) Representative example of the mtDNA structure from a sequencing read in one Petite sample. This plot displays a sequencing read wrapped around itself in a spiral. Coloured blocks represent annotated features in the reference genome which in this sample are two origins of replication that have been excised from the wild-type genome and stitched together into a concatemer structure. Red bars are the breakpoint locations. Black regions are unmapped near the ends of reads due to adapters and barcodes. c) Representative Grande read from a Grande colony showing a portion of a linear segment of the genome, without breakpoints present except at the ends of the reads (red bars) which mark the end of alignments. d) Summary of mtDNA breakpoints detected across 38 Petite colonies, that were derived from 9 spontaneous petite colonies through passaging (above diagonal), and 10 Grande colonies (below diagonal). In this scatter plot each marker represents the centroid of a cluster of mtDNA breakpoint signals in reference coordinates from reads in a single sample. The numbers of breakpoints in each family are indicated in the legend, as well as the numbers of colonies in each family sequenced. Also indicated are the regions on the reference genome that make up the repeating unit in the Petite read shown in (b) and the subsection of the reference genome that contributes to the Grande read in (c).

All Petite colonies contained at least one breakpoint signal and often a diverse set of breakpoints totaling 84 breakpoints across 38 Petite colonies, whereas in Grande colonies sequenced only 2 had high confidence breakpoints detected, with a total of 3 breakpoints across 10 Grande colonies (Figure 2d). The diversity in location of mtDNA breakpoints within Petite families and breakpoint counts greater than the number of members of each family/sub-family also echo the diversity observed in the coverage plots but with more detail; These diverse breakpoint distributions within families indicate either structural diversity in the progenitor colony, continued changes in mtDNA structure resulting in sub-families or coexisting structures in colonies, or multiple breakpoint signals within colonies indicating more complex mtDNA structures generated by multiple excision events.

#### Locations of excisions

The prevailing theory for the formation of Petites relies on sequence-specific illegitimate recombination within the wild-type DNA molecule between repeated GC clusters and AT stretches (Bernardi and Bernardi, 1980; de Zamaroczy et al., 1983; de Zamaroczy & Bernardi, 1986), which are prevalent in all non-coding regions of the mitochondrial genome in yeast. In particular, the extensive homology of the eight mitochondrial origins of replication and their inclusion of similar GC clusters (de Zamaroczy & Bernardi, 1986) suggest important regions for illegitimate recombinations. Evidence for hybrid origins resulting from recombination between adjacent origins in the wild-type genome have been seen in restriction digestion data (Marotta et al., 1982). Large structural variations and smaller mtDNA variations have also been observed in Illumina sequencing of Petites to cluster within origins and within close proximity to origins (Osman et al., 2015). Whether this theory of origin-origin recombination, or preferred excisions near replication origins holds up, we were curious to understand the role replication origins might play in the excision process. Placing non-inverted breakpoint locations and replication origin locations on mitochondrial reference coordinates reveals the clustering of breakpoints near edges of interacting origins of the same orientation (Figure 3a). In fact, ~30% of breakpoints reside within replication origins, indicating that the structures containing these breakpoints have origins that are perturbed by excisions, or hybrid origins. An additional ~20% of breakpoints reside within 275bp of the edge of an origin. The remaining 50% of breakpoints are located from 275bp to 3kbp from the edge of an origin.

**Figure 3:**
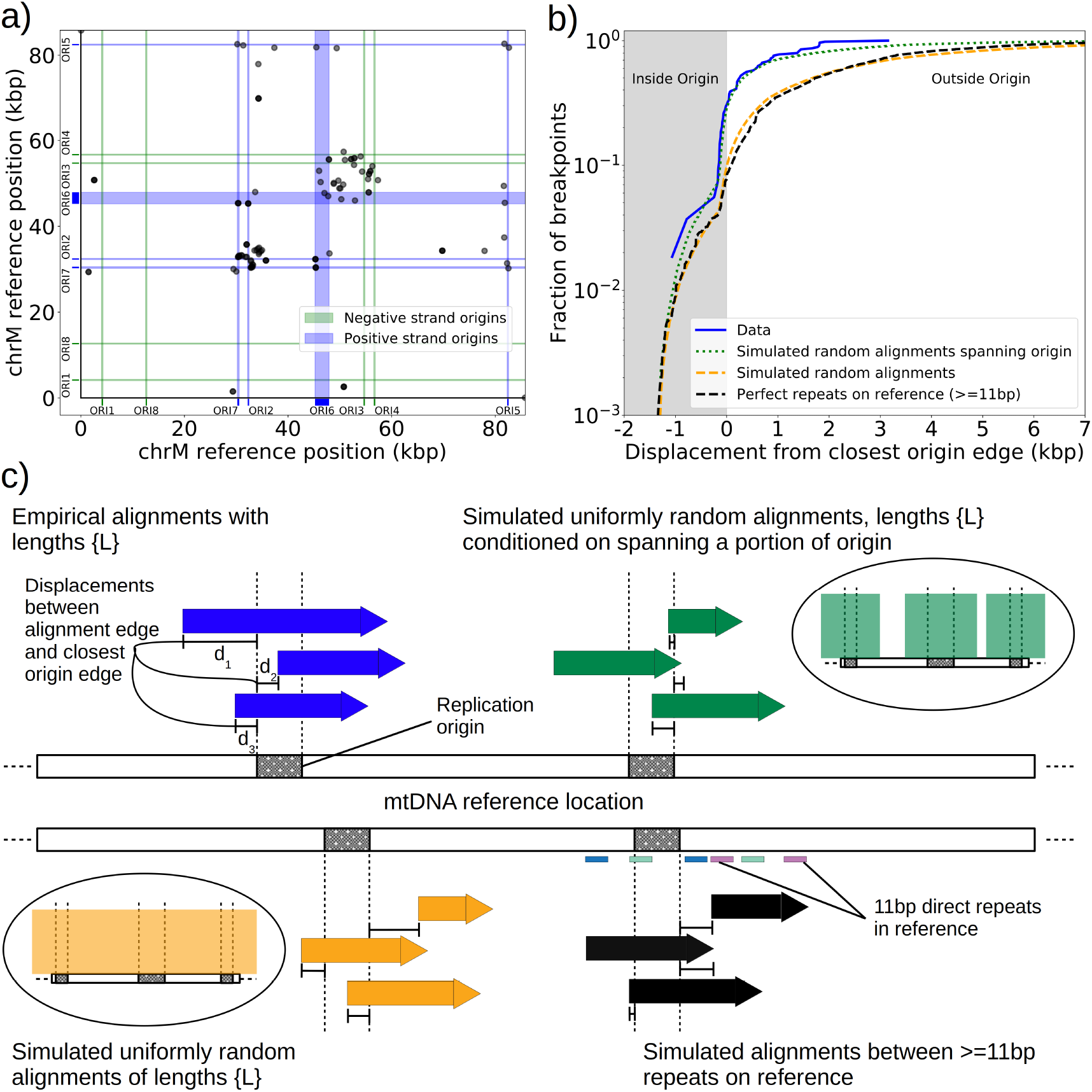
a) Black dots represent the centroids of breakpoint clusters (see methods), and blue and green shading highlights replication origins and their orientation. b) A cumulative plot of the displacements between breakpoint edges and closest origins of replication, where the blue curve shows this enrichment of breakpoints near replication origins (top left panel (c)). The orange curve represents a simulation of uniform random alignments placed on the reference genome following the true alignment length distribution in the data (bottom left panel (c)). The black curve represents the simulation of alignments between randomly selected perfect repeats >= 11bp on the reference sequence (bottom right panel (c)). The green curve agrees much better with the data (blue) curve, which is the same simulation of random alignments placed on the reference following the length distribution of the data, but with the requirement that these alignments span some portion of a randomly selected origin of replication (top right panel (c)).

Next, we asked if this non-uniform pattern of excision revealed new rules for mtDNA fragmentation. Besides the potential role of homology of the replication origins, it has been noted that a high density of unperturbed replication origins in Petite structures result in a replication advantage for Petite mtDNAs over wild-type mtDNAs (Blanc & Dujon, 1980; Zamaroczy et al., 1981; Mangin et al., 1983). In concatemer structures, this means that smaller repeated fragments containing replication origins are more fit than wild-type fragments when in competition with each other. This leads to a natural question of whether or not clustering near origins is due to higher frequency recombination within or near replication origins, or if it is due to selection on a pool of arbitrary excisions with selection for the resulting small fragments containing replication origins. To this end, we compared the distribution of displacements between breakpoints and the closest replication origins (Figure 3b, blue curve) to three different models. In the first model (Figure 3b, orange curve) we plotted the same displacement distribution for uniform random mtDNA fragments, with a size distribution given by the sequencing data, placed on the reference mitochondrial genome. In the second model (Figure 3b, black curve) we plotted the displacement distribution for random fragments between perfect repeats of greater than 11bp in the reference, which is motivated by the fact that excisions require perfect repeats or highly homologous regions. In the third model (Figure 3b, green curve) we plotted uniform random fragments as in the orange curve, with a length distribution from the sequencing data, but conditioned on spanning a portion of a randomly selected origin of replication. A schematic summarizing these models is provided in Figure 3c.

The orange and black distribution capture the breakpoint displacements from origins expected from random excision events and no selection for replication origin containing fragments. Note that the similarity between the black and orange curves demonstrates the prevalence of repeated homology in the mitochondrial reference genome. The green curve captures random excision, but strong selection for small replication origin containing fragments due to the requirement for alignments to contain a portion of an origin in this model. The observed data agrees nicely with the green curve for most of the domain of the distribution. The empirical origin to breakpoint distributions and their agreement with the green model are also consistent across a variety of breakpoint clustering/filtering parameter regimes, which have minor effects on the individual breakpoints extracted from the sequencing data, but little effect on these distributions (Table S2, Figure S2). Thus, while we cannot exclude a model of non-random excisions favouring close origin proximity, the bulk of the minimum breakpoint to origin displacement distributions observed can be explained by random excision and strong selection for small origin containing fragments which is in agreement with the prevailing theory of Petite mtDNA formation.

### The nature of structural diversity within sequenced samples - coexisting isoforms and resolution of heteroplasmy vs homoplasmy

#### Excision cascades, their patterns, and the structural diversity that results from them

Given the above described excision pattern and knowledge of the mechanism that generates Petite mtDNA from Grande mtDNA, we were curious to know if subsequent excisions in Petites differed from initial excisions in Grandes that generated the first Petite mtDNAs. In 16 Petite samples sequenced from families {1_a_, 1_b_, 2, 3, 8_a_} there were detectable levels of repeated structures that differed from the “primary” mtDNA structures which span the longest portion of the reference genome and generally contribute the majority of mtDNA (see methods on details of structure reconstruction). These lower frequency structures, or “alternate” structures as they will be described from here on, were found to contribute from 0.1% to 59% of total mitochondrial content in these samples. Following multiple passagings of Petite colonies before sequencing, which would rapidly dilute any initially coexisting structures due to mtDNA bottlenecks (Ling & Shibata, 2004), these alternate structures most likely result from subsequent excisions of the primary structure during culturing. Such “excision cascades”, where further excisions act on existing Petite fragments were hypothesized and discussed by (Locker et al., 1979; Marotta et al., 1982; Bernardi, 2005), where it was suggested that the varying levels of alternate structures will depend on their generation rate and selective advantage in replication over the primary structure. Part of what makes the dynamics of mtDNA within these cascades interesting is the multiplicity afforded by a large variety and number of potential excisions; Particular excisions that bring regions of homology together may open up entirely new trajectories of excision dynamics that were previously unlikely or inaccessible due to mtDNA conformation.

An extreme example of such an excision cascade is given in Figure 4a, where both primary and alternate alignments are shown on the reference mitochondrial genome location in addition to sequencing coverage. In Figure 4b the structures of the repeated units composed of these alignments are also shown alongside their calculated mitochondrial content frequencies (see methods - Primary/alternate structural frequency calculations). The first thing to note is that the locations of alignments extracted from our structural detection pipeline that are involved in repeat units align well with the total sequencing coverage in Figure 4a. This diversity of alignments in Figure 4a is also corroborated by colonies that share the same progenitor; Each of the coverage curves of members of family 1 in Figure 1b share peaks with the coverage curve in Figure 4a. Secondly, there is significant diversity in the type of repeat units and necessary steps in their generation which stitch together these alignments in Figure 4b: Repeat unit #1 shown in Figure 4b (yellow and pink) is an example of a secondary excision across a segment of mtDNA containing the preexisting primary breakpoint, as both alignments come from opposite ends of the primary alignment and are stitched together. This immediately suggests that the excision occurred in mtDNA in a concatemer form, and across the repeat unit breakpoint (green and purple Type II excision, Figure 4c). Repeat unit #2 (blue, maroon, green) also spans the primary breakpoint, but has an additional alignment in a different orientation that either resulted from two excisions or recombination of different repeat units (green, purple, and gray Type II excision, Figure 4c). Repeat unit #3 (green) is an example of an excision within the primary repeat unit and away from its edges (Type I excision, Figure 4c). Repeat unit #4 (purple) is an example of a repeat unit that shares only one edge with the primary alignment, but with this edge interacting with a different region of the genome producing a new alignment (Type III excision, Figure 4c).

**Figure 4:**
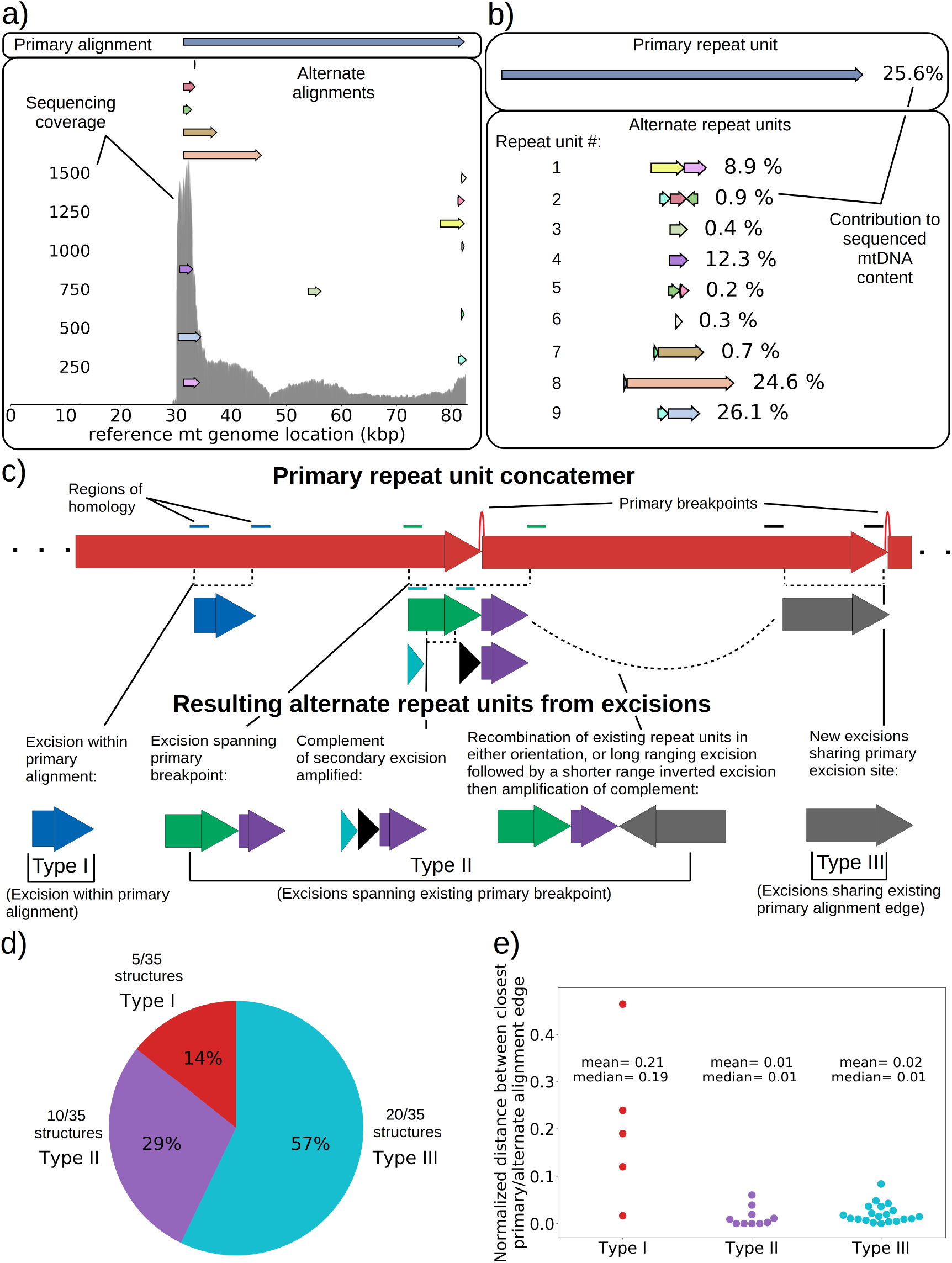
a) Example locations of alignments from mapped reads (linear alignments here are bounded by breakpoints) observed in long reads within Petite sample 1b. Endpoints of arrows are the mean breakpoint location for the cluster of breakpoint signals that punctuate alignments in repeats. The first panel shows a primary alignment which has the longest span on the genome and exists in long repeats. The second panel shows smaller alternate alignments that exist within detected repeated structures at a lower frequency. Also included in gray is a sequencing coverage map of this sample. b) Excision cascade in Petite sample 1b. This plot shows the same primary repeat unit in the first panel, and its contribution to total mitochondrial content as a percentage. The second panel shows the forms of alternate structures present in the same sample which were derived from the primary alignment, alongside their mitochondrial contribution as a percentage. c) A schematic of the multiplicity of excision events that generate alternate repeat units. The primary concatemer is the red structure, where arrows indicate the alignments (contiguous regions of the reference) that make up the repeating units. The primary breakpoint between these alignments is denoted as a red arc. Coloured rectangles above the alignments with the same colours indicate regions of homology in the primary structure that can interact to produce an excision. Dotted rectangles indicate excision sites that produce the alignments shown below them. In the lower half of the figure, five distinct excision events that can generate different repeat units are shown. These are grouped into three excision classes in the data: Type I, where excision occurs within primary alignments, Type II, where excisions span the existing primary breakpoint, and Type III, where excisions share one edge of the primary breakpoint. d) The frequency of each class of excision across 35 alternate structures detected in the data. e) A plot of the distance between alternate alignment edges and their closest primary alignment edge across all three classes, normalized by primary alignment length.

In Figure 4c we provide a schematic of the types of alternate repeat units observed across all samples and the plausible mechanisms of generation. We classify the resulting alternate repeat unit into three easily distinguishable classes in our data: Type I alternate repeat units are regions excised from the interior of primary repeat units. Type II alternate repeats contain or span the primary breakpoint, resulting from an excision across the breakpoint between two primary repeat units in a concatemer form. Type III alternate repeat units share one edge with the primary breakpoint and have a new edge within the primary alignment. For the technical details in the classification of these repeat unit types, see methods - TypeI/II/III repeat unit classification. The proportions of each class of alternate repeats (35 total) across all samples are shown in Figure 4d, where it is clear Type III breakpoints make up the majority (57%) of alternate repeated structures observed across the 16 colonies where we see alternate structures. In Figure 4e we also plot the distance between the closest primary and alignment edges for each class of repeat normalized by the primary alignment length. It is clear from this figure how close subsequent excisions are to primary breakpoints in the most abundant class, Type III, with a mean and median fractional distance between the alternate and primary edge of 2% and 1%, respectively. The mean and median fractional distances of Type II and Type III repeats are also comparable, as expected for structures that share edges of the preexisting primary breakpoint. The abundance of Type III repeats suggests a strong preference for secondary excisions at the site that produced the primary alignment itself, that ultimately constrains the trajectories of subsequent excision events in an unexpected, and previously unreported way. It is unclear at this time whether Type II repeats, which encompass intact primary breakpoints, are related to this phenomenon. In general, this pattern of a preference of excisions around or across primary breakpoints is consistent across a variety of clustering/filtering parameters for breakpoints detection (Table S1-S2, Figure S3), the details of which are described in methods.

#### Heteroplasmy in “mixed” structure examples - Coexisting isoforms and intramolecular heterogeneity

Seven colonies within family 3 in Figure 1b displayed distinguishably higher variance in coverage than the rest of the Petite colonies sequenced. This variance in coverage suggested either a complex repeat unit which itself contained smaller repeated units, or heterogeneity of mtDNA content in these samples. As such, we were interested to understand the source of this coverage variability. In these colonies, sequencing revealed non-periodic primary structures involving partial inverted duplications of sequences. This is in contrast to the repeated units as concatemers that are found in the remainder of the Petite colonies and are primarily in tandemly repeated (non-inverted) forms. These non-periodic structures resemble the “mixed” structures first characterized in detail in (Heyting C, et al. 1979). The structure of one of these colonies is detailed in Figure 5, and is representative of all 7 “mixed” structure colonies as they contain indistinguishable alignments and were derived from the same spontaneous Petite colony. In Figure 5a, four alignments of different lengths and reference locations are depicted as arrows. Note that all four alignments share the common region of the red alignment. Also included is the sequencing coverage of this particular colony, which aligns nicely with these alignments extracted from our structural repeat detection pipeline, as well as the ranked length and absolute count of alignments. Correcting for sampling bias (see methods - Mixed structure alignment frequency calculations) due to the sampled read length distributions across 7 colonies with this same structure reveals that each alignment exists in equal proportions in colonies that harbour this structure (Figure 5b). Example structures in long reads selected from one of these samples are provided in Figure 5c, where the mixed structure is evident with seemingly random orientations of alignments. These “mixed” structures are clear examples of intramolecular heterogeneity in mtDNA and likely intermolecular heterogeneity across a population of mtDNA fragments within cells given the differences in content of the fragments observed. To attempt to make sense of this structure, which is at odds with the concatemer structures observed in all other colonies, we applied the repeat detection pipeline to see if any reads exhibited repeated structures. Interestingly, while superficially this structure seems devoid of a clear pattern and appears uniformly randomized, across all 7 colonies with the structure shown in Figure 5c we did see some evidence of partially repeated structures, where the same alignments were repeated with the same orientation but separated by other single inverted alignments (Figure 5d). In all 7 colonies with these same four alignments, these partially repeated structures are composed of the concatenation of largest and smallest alignments with opposite orientation, or the concatenation of the second-largest and second-smallest alignments.

**Figure 5:**
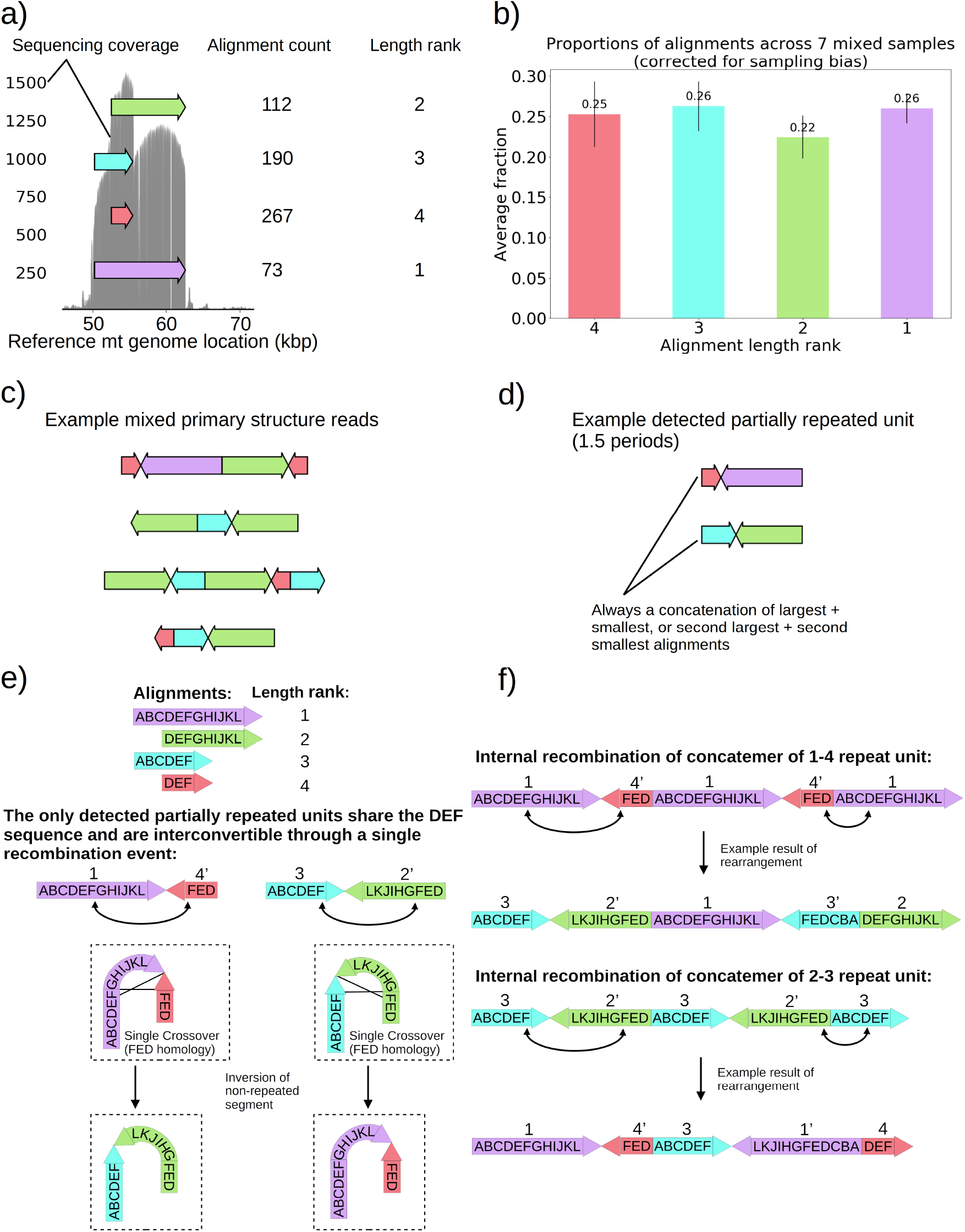
a) Alignment locations and their raw counts present in the primary structure of a “mixed” repeat Petite colony. b) Proportions of each four alignments across 7 mixed petite colonies sequenced after accounting for sampling bias (see methods). c) Example structures in sequencing reads in this colony, displaying a collection of coexisting isoforms with identical base pair content but varying structures with two distinct inverted duplication breakpoints delimiting alignments. d) Example partially repeated units detected after observing 1.5 periods in the repeat detection pipeline. e) Interconvertibility of detected partial repeats. Arrow directions indicate the strand to which alignments have been mapped, in addition to the prime notation on length ranks of each structure which indicates an inverted alignment. f) Crossover events in the background of concatemers that can produce all breakpoint transitions and structures observed in the data.

The detection of the partial repeats in Figure 5d and the overlapping context in Figure 5a led us to the proposed mechanism of generation provided in Figure 5e-f, which with this new evidence builds upon on the crossover mechanism first hypothesized in (Bos et al., 1980) for what we believe is the same structure we observed. In Figure 5e we show how the only detected partially repeat units are interconvertible through a single crossover event relying on interactions of oppositely oriented regions. This crossover event in concatemers or repeat units results in the inversion of the non-repeated sequences, and such an event can produce all of the “mixed” read structures we observed across 7 samples (Figure 5f). The generative picture given this proposed mechanism is the following: (1) First a recombination event produces one of the repeat units in Figure 5d, by a crossover mechanism like that suggested in (Faugeron-Fonty et al., 1983), or origin dependent mechanisms like those observed in yeast and proposed in (Brewer et al., 2011; Brewer et al., 2015). (2) This repeat unit is amplified, forming a concatemer through rolling circle replication, which exists in this form only transiently. (3) High frequency recombination at the region of shared context which was also suggested by (Bos et al., 1980) produces nearly uniformly random orientations of alignments in a concatemer form. This proposed mechanism, and the fact that 7 colonies derived from the passaging of one spontaneous Petite all had this “mixed” structure, strongly suggests that cells in these colonies are heteroplasmic in these various structures because they are not readily segregated. As such, this structure represents a unique example of coexisting structural isoforms in the mtDNA of baker’s yeast, that are produced through rapid recombination events that counteract the periodic structures produced by rolling-circle replication and any strong selection for particular configurations of mtDNA.

#### Evidence of both Heteroplasmy and Homoplasmy within Grande and Petite colonies

Given the evidence of heteroplasmy in “mixed” structure colonies, we were curious to understand the nature of the low frequency alternate mtDNA structures we observed in both Grande and Petite colonies. In the bulk sequencing of colonies, low frequency structures can be contributed by both heteroplasmic cells and mixed populations of cells homoplasmic for primary and alternate structures. In the heteroplasmic limit (Figure 6a), the majority of alternate structure content in a colony is contributed by cells in a heteroplasmic state. One consequence of being in this limit is that biological replicates of colonies would be expected to have low variance in total alternate structure content if heteroplasmy persists. In the homoplasmic limit, the major contribution to total alternate structure content comes from cells solely containing alternate structures (Figure 6b). In this limit, stochasticity in the time of generation of the mutational event would be expected to result in high variance in alternate structure content across biological replicates. Furthermore, in circumstances that enable differential selection on homoplasmic lineages, such as in Petite lineages within Grande colonies in non-fermentable conditions, one would expect the alternate structure content to change as a function of growth conditions if there were homoplasmic contributions.

**Figure 6:**
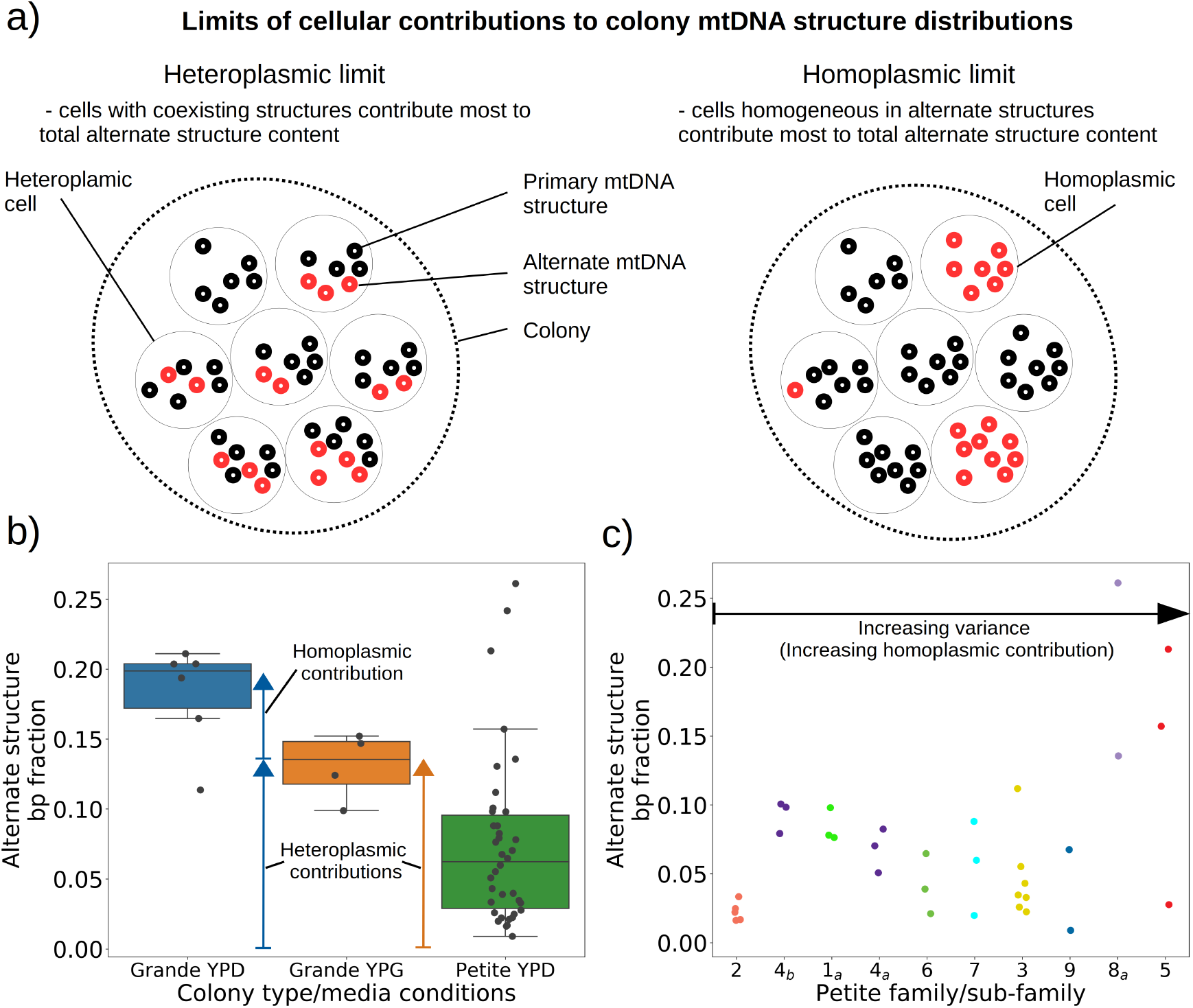
a) A schematic of the two limits of cellular contributions to mtDNA colony structure. Left: In the heteroplasmic limit, most of the contribution to total alternate structure content comes from cells containing coexisting alternate and primary structures. Right: In the homoplasmic limit, most alternate structure content comes from cells homogeneous in alternate structure content. b) The total fraction in bp of reads that include any breakpoint not expected from the primary structure in Grande samples in YPD (fermentable carbon source), YPG (non-fermentable), and Petite samples in YPD. Each dot represents the alternate structure content fraction for a single colony, which is the fractional contribution to total mitochondrial content of reads that contain breakpoints that differ from breakpoints in the primary structure. The box plot displays the median value, and the minimum, maximum, 1st quartile, and 3rd quartile. Blue vectors indicate heteroplasmic/homoplasmic contributions in Grande colonies in YPD. The orange vector indicates heteroplasmic contributions in Grande colonies in YPG. c) Contributions of petite families/subfamilies to the Petite YPD alternate structure bp fractions in (b) sorted by variance in alternate structure basepair fractions in each subfamily. The arrow indicates that increasing variance is expected to be accompanied by increasing homoplasmic contributions.

In 10 Grande colonies sequenced, 4 were grown in YPG (non-fermentable media) and 6 in YPD (fermentable media). Only one colony in each growth condition harboured high confidence Petite concatemer structures according to our structural detection pipeline, represented by the two distinct breakpoint clusters in the lower half of Figure 2d. One of these high confidence structures within a YPG colony is shown in Figure S4. With a spontaneous Petite frequency of 10% in the genetic background of the strains sequenced (Dimitrov et al., 2009), these low detection rates are due to our conservative approach to detecting breakpoints; In our pipeline we require at least 3 breakpoints from three separate reads to form a believable cluster of breakpoint signals (see methods). In Grande colonies that produce a diverse set of Petite structures afforded by excisions of the intact WT genome, forming high confidence breakpoint clusters, let alone clusters themselves, is unlikely. Therefore, to compute alternate structure frequencies in this analysis we abandoned the requirement of breakpoint signals to form a cluster. Instead, we simply counted the total base-pair contribution of reads that included any detected breakpoints internal to the primary alignments as long as they were not accompanied by inverted duplication artifacts which are known to introduce spurious breakpoints due to noise in the latter half of the read (Spealman et al., 2020. and Figure S5).

The results of this read enumeration approach (Figure 6b) indicate that Grande colonies grown in YPD (fermentable media) have a higher mutant mtDNA frequency than those grown in YPG (non-fermentable media). Knowing that Petite cell lineages experience strong negative selection in YPG, these results suggest that this elevated alternate structure frequency in Grande YPD colonies is due to clonal divergence, resulting in homoplasmic Petite lineages that contribute at least in part to the observed signal. At the same time, the presence of alternate structures in YPG media also indicate heteroplasmic contributions (orange vector in Figure 6b), as these alternate structures are unable to exist in homoplasmic lineages for long times under non-fermentable conditions and will be rapidly diluted during growth. Thus, these results in Grande colonies demonstrate that both heteroplasmic and homoplasmic cells (sum of blue vectors in Figure 6b) are contributing to alternate structures in fermentable conditions.

In Petite colonies which only grow under fermentable media conditions, cells regardless of their mtDNA content have similar fitness. Therefore, in Petite colonies clonal divergence goes unchecked, which would be expected to result in a high variance alternate structure fraction within Petite colonies. This is indeed what is seen within Petite colonies in Figure 6b. It is also clear that the median alternate structure frequencies in Petites exist below those of Grande colonies in YPG. This we suggest is due to the out competition of alternate structures by primary structures that are more fit in Petites than in Grandes, indicating at least transient heteroplasmy. To further explore where the variance in alternate structure frequencies comes from in Petite colonies, we grouped points in Figure 6b by the Petite families/subfamilies that contributed them, sorted by variance (Figure 6c). Given the presence of genetic bottlenecks and within-cell selection, the majority of the variance in alternate frequency within families is expected to be due to stochasticity in the generation of Petite lineages homoplasmic for alternate structures. However, we do observe a large spectrum of variance in Figure 6c, with some families on the left of the x-axis such as 2 and 4b having distinguishably lower variance in alternate structure frequency than the families at the rightmost limit of the x-axis, such as family 5. The low variance in the leftmost families on the x-axis suggest that heteroplasmic cells may contribute significantly to their total alternate structure frequencies. Thus, as in Grande colonies, we are able to tease apart indications of heteroplasmic and homoplasmic contributions in Petites.

### Structure-function relationships of mtDNA in yeast

#### The role of origins of replication and GC clusters

The mechanism of generation of Petite mtDNAs, as well as our explanation of the distribution of excisions observed in spontaneous Petites, relied on the presence of replication origins. Given their importance in conferring a replication advantage to Petite mtDNAs, we were interested to look for mtDNAs without replication origins that we expected would exist at a low frequency. We were also curious to know if mtDNAs devoid of replication origins had any shared structural characteristics that might explain their propagation. Consistent with the notion that repeated structures with high densities of replication origins have a selective replication advantage over wild-type Grande mtDNA (Zamaroczy et al., 1981; Bernardi, 2005), 32 of 47 repeat structures detected across the 16 Petite colonies that contain alternate structures exhibit a higher replication origin content fraction than wild-type mtDNA (Figure 7a). Furthermore, all primary structures contain at least a portion of an origin. However, some of the detected alternate structures encircled in red in Figure 7a have no replication origin content at all. These resemble the “surrogate” replication origin structures described in (Goursot et al., 1982) and appear to contain GC clusters in similar configurations to replication origins (peaks above 0.6 GC content in Figure S7), which are known to be important for replication and transcription initiation (Baldacci & Bernardi, 1982; de Zamaroczy & Bernardi, 1986). Consistent with this idea, 7 of 9 structures without replication origins are enriched in GC clusters compared to the average GC cluster content of wild-type mtDNA (Figure 7b). Besides the suggested involvement of GC clusters in replication, the enrichment in GC clusters here is also consistent with the observation that GC clusters themselves may be preferred over AT rich regions as excision sites (Faugeron-Fonty et al., 1979; Gaillard et al., 1980). While structures without replication origins are rare in cultured spontaneous petites (Goursot et al., 1982), high depth long read sequencing has provided access to these low frequency structures. The ease of identification of these mtDNA structures through long-read sequencing and accompanying structural inference techniques may prove useful in exploring the minimal sequences required for replication in yeast, as well as low frequency genome diversity in other systems.

**Figure 7:**
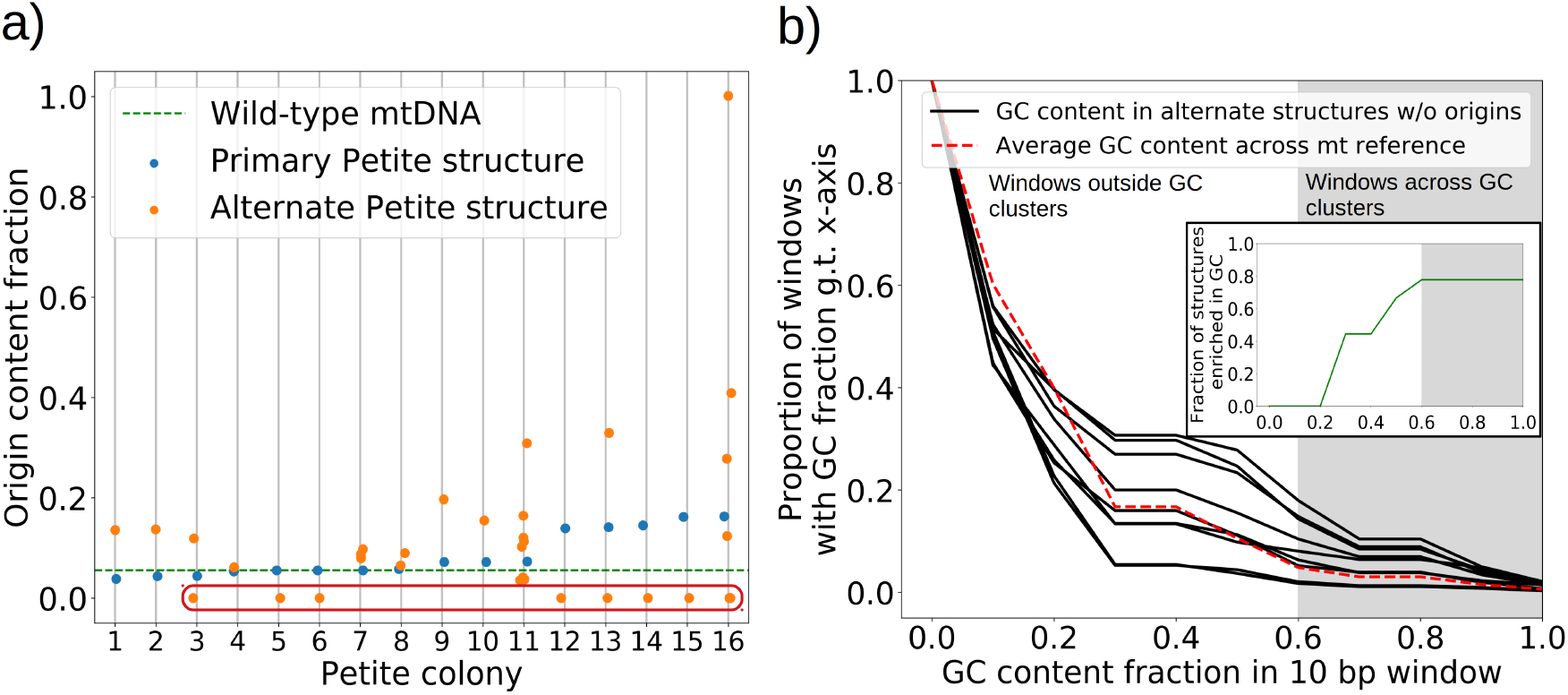
a) Replication origin content fractions in primary/alternate structures detected in all samples where both are present. Each dot represents the base pair fraction of any of the eight origins of replication in detected structures. Orange dots are the origin fractions in alternate structures, blue in primary structures, and the green line is the origin content fraction in the wild-type mitochondrial genome. Highlighted by a red bubble are nine alternate structures that are devoid of an origin of replication. b) Black curves (9 total) represent the cumulative distribution of GC content fraction in a sliding window of 10bp in the highlighted zero-ori alternate samples. The red curve highlights this same GC distribution but in the wild-type (Grande) mitochondrial reference. The gray region indicates GC content fractions in sliding windows that are consistent with GC clusters found in replication origins (Figure S7). The inset shows the fraction of black lines above the red line as a function of GC fraction in the 10bp window.

#### How mtDNA structure informs suppressivity

To understand the rules of competition between wild-type and mutant mtDNA, we measured the suppressivity of all Petite colonies within families (see methods). Suppressivity is a measure of the fraction of Petite progeny in a cross between each Petite sample and a Grande tester strain. Unlike previous work that studied the relationship between structure and suppressivity in highly suppressive Petites with suppressivity upwards of 90% (de Zamaroczy et al., 1981), our strains exhibit suppressivities from the basal rate of the Petite frequency of the Grande strain at 10%, to ~90%, and these suppressivities correlate well with repeat unit lengths of up to 70kbp (Figure S8). In contrast, the repeat units in previous work were smaller than 10kbp. While this difference in repeat unit size was due to intentional selection of small repeat units in the previous work, distributions of deletion sizes and therefore observed suppressivities have been shown to be dependent on numerous nuclear genes (Bradshaw et al., 2017; Ling et al., 2019). To describe how the structure of mtDNA in our samples informs suppressivity, we developed a phenomenological model (Figure 8a) which assumes each repeat unit is independently competing (we discuss alternate models in Figure S9). The key assumptions of the model that explains the data well are: (a) in mating both Grande and Petite cells contribute equal mitochondrial content, *M*, which is motivated by the observation of equal Grande and Petite contributions observed in (MacAlpine et al., 2001), (b) the number of repeat units initially contributed by Grande and Petite cells during mating is given by *M* over the average repeat unit length (*L_G_* or *L_P_ G* for Grande, *P* for Petite), and (c) Petite and Grande repeat units replicate independently and exponentially with a replication rate linearly dependent on a mtDNA replication speed (*ν_G_, ν_P_*) and replication origin density (*ρ_G_, ρ_P_*). The suppressivity is then the fraction of Petite repeat units after a certain competition time (*t*) and is given by the ratio of the time evolution of an exponentially growing population of Petite repeat units *N_P_* = (*M/L_P_*)*e^νPρPt^* to total repeat units *N_P_* + *N_G_* = (*M/L_P_*)*e^νPρPt^* + (*M/L_G_*)*e^νGρGt^*, given in equation (1):

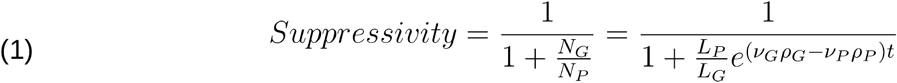

**Figure 8:**
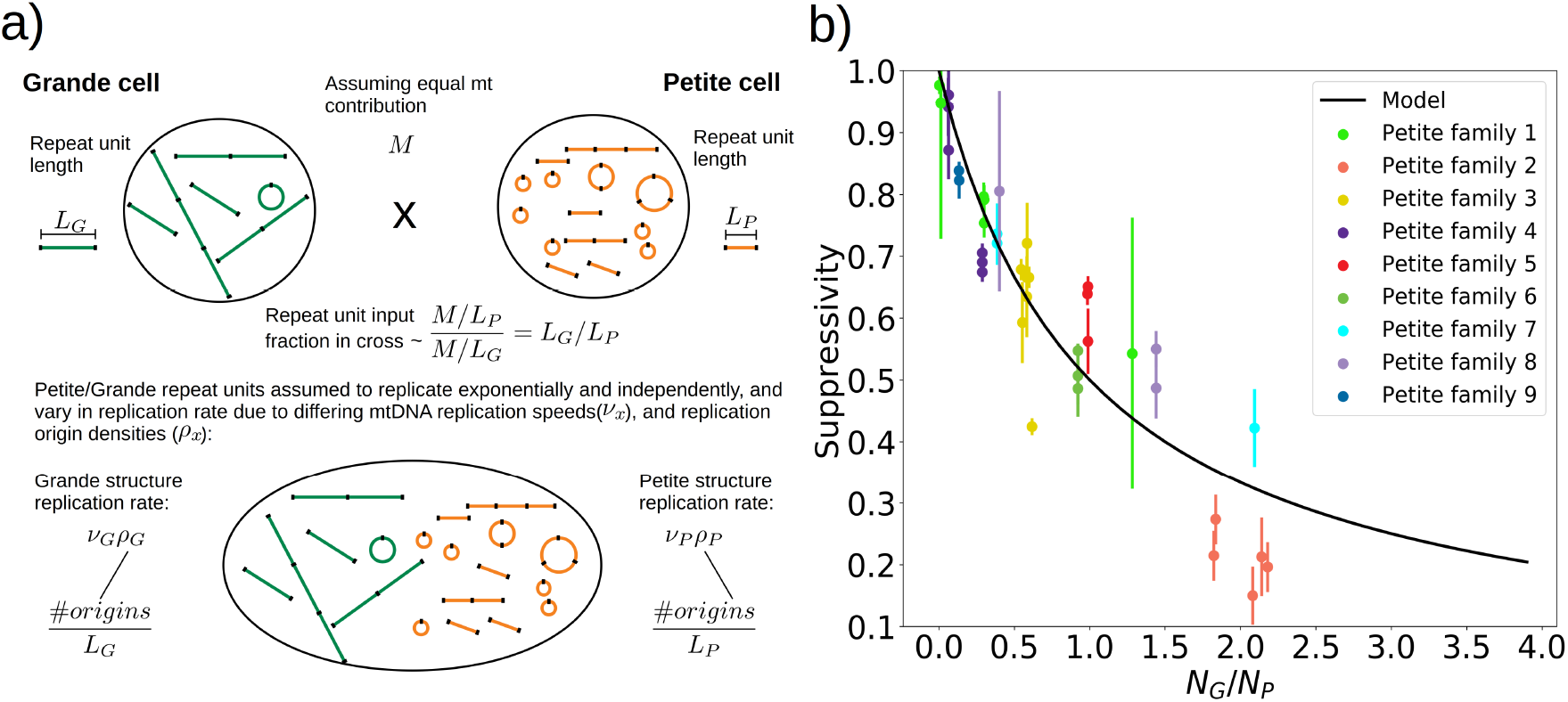
a) A visual depiction of a phenomenological model of suppressivity. Grande and Petite cells are assumed to contribute equal quantities of mtDNA. It is also assumed that each repeat unit replicates independently and exponentially and that during mating the repeat unit input fractions of Grandes and Petites are inversely proportional to repeat unit length. Exponential growth rates are the product of mtDNA replication speeds and origin densities. b) Suppressivity of all samples compared to a fit of equation (1), which is the black line. The fit parameters are *ν_g_t*\* = 10677, *ν_p_t*\* = 2296;, and the coefficient of determination is *R*^2^ = 0.85. Dots are the average suppressivity across three second passage Petite colonies that share the same first passage progenitor, and belong to the families indicated in the legend (same coloured dots share a spontaneous Petite colony progenitor). Y-axis error bars are +-the standard deviation in suppressivities across these three second passage Petites colonies. Samples containing inverted breakpoints in their primary structure are those derived from family 2 and 3, the orange and yellow dots, respectively. Family 3 are the mixed structures described in the maintext.

The data and least squares fit of this model, with *ν_G_t*\* and *ν_P_t*\* as fit parameters, are shown in Figure 8b. These fit parameters are the products of mtDNA replication speeds in Grandes and Petites with the competition time (*t*\*) over which competition of Grande/Petite structures occurs following mating. The repeat unit lengths *L_G_* and *L_P_* in equation (1) are taken to be the sum of unique alignment lengths in each sample, and *L_G_*/*L_P_* is the second term in the denominator of equation (1), which is the ratio of the Grande to Petite fragment population. To understand the values of the fit parameters and whether or not they are reasonable, we compare them to equivalent parameters inferred from an exponential growth model of a budding Grande cell population: First we assume that the competition window (*t*\*) is equal to the doubling time in a diploid Grande population of cells (90 minutes). If we also assume that the exponential replication rate of the budding population is the product of replication origin density (1 every 10 kbp in Grandes) and replication speed, then the average mtDNA replication speed in Grande cells is 82 bp/min. This is of the same order as both of the *ν_G_* and *ν_P_* fit parameters in Figure 8b assuming that the supressivity data is a result of competition over the same doubling time.

With respect to the architecture of the model, a variety of alternative models were also tested in Figure S9, revealing that both exponential growth and a repeat unit input fraction inversely proportional to repeat unit length are statistically important inclusions in improving the model in most regimes. The inverse repeat unit length terms seem to suggest that within yeast zygotes, early competition operates in a repeat unit limit where concatemers are reduced to monomeric forms which then undergo replication. Interestingly, active concatemer to monomer partitioning has been observed during mitosis in yeast (Ling & Shibata, 2002, 2004, 2007), although to our knowledge little is known about the structure of mtDNA during mating and zygote formation. Thus, according to this model, the rules of competition between wild-type and mutant mtDNA in yeast depend on the exponential replication of monomeric forms of mtDNA in zygotes, where replication rates are proportional to replication origin densities in repeat units. This highlights the possibility that Petite mtDNAs may have both a replication advantage and segregational advantage if replication occurs in physically separated repeat units in zygotes.

## Discussion

Petite mtDNA in yeast has been extensively studied with early sequencing technologies since the discovery of the Petite phenotype in (Ephrussi, 1949; 1953). This was a system of interest because of the dispensability of mtDNA, which allowed exploration of destructive mutational events and because of abundant non-coding sequences which at the time had no known function. These studies revealed the structure and role of mtDNA replication origins, the excision mechanism that generated Petite mtDNAs, and explanations for the replication advantage of Petite mtDNA over wild-type mtDNA that are responsible for their existence. In this article we studied mitochondrial genome dynamics of Petite mtDNA through single-molecule long read sequencing of spontaneous Petite colonies. This sequencing technology, in conjunction with structural inference methods we developed, which to our knowledge have never been applied to Petite mtDNA in yeast, gave us the ability to reconstruct complex mtDNA structures with high resolution within populations of growing Petite colonies. Among results echoing studies spanning the entire history of research of this genome in *Saccharomyces cerevisiae*, we showcased a previously unseen pattern that constrains subsequent excision events in generating new Petite structures from existing ones, settled contention in the literature surrounding the existence and generation of non-periodic “mixed” Petite structures, and proposed a phenomenological model of suppressivity.

First, we demonstrated that the colocalization of Petite inducing mtDNA excisions and origins of replication could be explained by selection for small origin-containing fragments and not necessarily preferential excisions near origins. The origin-excision proximity we observed is consistent with (Osman et al., 2015). However, the origin selection explanation provided here calls into question the hypothesis put forth in (Osman et al., 2015) which is that the origins of replication are mutational hotspots for both small variations like SNPs and larger structural variations. On the other hand, rare single-base changes at inferred excision sites in Petite strains have been observed (Zamaroczy et al., 1983), suggesting that small mtDNA variations may be present at the excision sites that are clustered near origins due to selection, rather than within origins. As it stands, more work needs to be done to determine if excision sites in Petites are accompanied by small variations, or if smaller variations that we are unable to access in Nanopore sequencing are truly enriched within origins of replication. It is also important to note that the mtDNA recombination landscape in wild-type yeast (Fritsch et al., 2014) also varies significantly from the excision distribution we observed. However, we attribute these differences to the selection of origin containing fragments in Petites, as well as differences between homologous recombination (between DNA molecules) and sequence-specific illegitimate recombination (within individual DNA molecules) responsible for excisions.

Next we explored the evolution of mtDNA structure in excision cascades, providing an example in a strain with significant structural diversity, highlighting how we could be quantitative in computing structural frequencies directly from reads, and revealing an apparent preference of excision at or around existing excision sites. Various levels of heterogeneity within strains indicated by non-primary sub-stoichiometric bands in the restriction digests of Petite mtDNA have been commented on previously (Bernardi et al., 1976; Lewin et al., 1978; Lewin et al., 1979; Locker et al., 1979; Marotta et al., 1982). The presence of these low frequency structures suggest that the excision mechanism is ongoing, continuously producing lower complexity Petite structures in a hypothesized excision “cascade” (Locker et al., 1979; Bernardi, 2005). Here we have demonstrated unequivocal evidence of secondary excisions that operate on primary structures, and critically, their relationship to the primary structure in this excision cascade. Single-molecule sequencing enabled us to quantify the abundance of these structures within colonies and the inference of rare structures, many of which would be inaccessible upon attempted segregation in passaging due to their low generation frequency. By classifying excisions into three classes across all samples sequenced, we showed how an apparent preference for the reuse of existing excision locations or regions around them constrained the fate of structures formed through subsequent excisions. The reuse of excision sites highlights a tension between contingency and repeatability in the formation of new Petite mtDNA structures. It also seems to suggest that the breakpoints in primary Petite structures are persistent instabilities in mtDNA, perhaps akin to structures like R-loops (Holt, I., 2019) that may promote strand invasion and recombination at or near these sites.

Here we also provide direct evidence for intramolecular and intermolecular heterogeneity in reporting “mixed” non-periodic structures in the mtDNA of yeast. Hints of the structures that we are describing as “mixed” structures have been commented on previously in ethidium bromide treated Petites (Locker et al., 1974; Locker et al., 1979) and spontaneous Petites (Heyting et al., 1979; Bos et al., 1980; Faugeron-Fonty et al., 1983). The first proposal of a model for the generation of these structures was provided in (Bos et al., 1980), but was subsequently refuted in (Faugeron-Fonty et al., 1983) where the claim was that these were larger ranging periodic structures produced by an unknown mechanism. However, our evidence of partially repeated units in Figure 5d, where two alignments are repeated with the same orientation but separated by an inverted alignment, precludes the structure proposed in (Faugeron-Fonty et al., 1983) and is evidence that adds to and is consistent with (Bos et al., 1980). As such, evidence presented here suggests the model presented in Figure 5e-f, based on the initial hypothesis in (Bos et al., 1980) is the appropriate model of generation of these “mixed” structures and that they are truly non-periodic. Thus, this structure is indicative of rapid recombination within mtDNA concatemers that opposes the homogeneity produced by rolling-circle replication and segregation through bud bottlenecks, which results in a collection of coexisting structural isoforms of mtDNA. While coexisting concatemers of various lengths and forms have been observed in yeast mtDNA with the same repeat units (Locker et al., 1979), as well as coexisting isoforms in plant mitochondria (Kozik et al., 2019), the “mixed” structures we have observed are a rare glimpse of this phenomenon in yeast that provide a unique example of persistent structural heteroplasmy. These structures are also interesting from the perspective of reverse excision events thought to partition concatemers into monomers during bud formation (Ling & Shibata, 2002). It is unclear how a monomer should be defined in these “mixed” samples, and therefore how it is partitioned, given that circular repeat units are the predominant species in new buds.

On the nature of structural heterogeneity in Petite samples, we argued that most of this heterogeneity we observe is likely due to homoplasmic clones, but with hints of heteroplasmic contributions in a few examples. The persistent heterogeneity in Petite samples observed previously as sub-molar restriction digestion fragments (Lewin et al., 1979) is largely consistent with our results. These fragments were persistent across biological replicates, but seen to disappear and reappear with varying intensities during subcloning, indicating heteroplasmy due to their persistence and clonal divergence explaining their varying intensities. The advantage however, of the single-molecule approach we took is clear here, as we have been able to both quantify this heterogeneity and infer these alternate structures with confidence, which would normally require segregation and isolation of these low frequency Petite clones. In addition, the observation of alternate structures in Grande samples under non-fermentable growth conditions is also a strong indicator of at least transient heteroplasmy which is usually difficult to resolve from coexisting homoplasmic clones in bulk sequencing data. Sequence inference also gave us access to low frequency alternate structures that are devoid of replication origins, which turned out to be wonderful examples of non-coding “selfish” mtDNA that is replicated solely due to its replication function and involvement in replication initiation. These are regions of mtDNA outside canonical replication origins that in addition to origins likely confer a replication advantage to wild-type fragments containing them. While these structures are rarely isolated as primary structures in Petite colonies due to out-competition by parental genomes (Goursot et al., 1982), single molecule sequencing and structure inference has provided easy access to these reduced structures and may be of use in exploring low frequency structural diversity in other systems.

Using what we learned about inferring fine structure from long reads, we then explored how this structure informed the suppressivity of the Petite samples sequenced and developed a phenomenological model of suppressivity. The relationship between suppressivity and mtDNA structure has been explored in (Zamaroczy et al., 1981), which provided two general rules of how mtDNA structure informs suppressivity: (1) Partial deletions or rearrangements of origins of replication, including inversions of fragments containing origins, reduce suppressivity, and (2), suppressivity is inversely proportional to repeat unit length. This was followed with an observed exception to the second rule (Rayko et al., 1988), that suggested that flanking regions also had an effect on suppressivity holding repeat unit length and origin content relatively constant. Our results echo the predictive value of the repeat unit size. In fact, among the models explored the most predictive is in the repeat unit limit which assumes that both Grande and Petite mtDNAs are independently replicating in their monomeric forms. While the assumption of equal mtDNA contributions by mating cells in the model is largely uncontested (MacAlpine et al., 2001), the size distribution of concatemers between Grandes and Petites reported have been highly variable; Petites have been found to have both larger and smaller average molecular sizes than Grandes depending on the strain and in both cases harbour a pool of concatemers of various sizes (Locker et al., 1979). However, these observations were in haploid strains not during zygote formation and it is possible for a distribution of concatemer lengths with a small average size to be consistent with the most predictive model explored in this work.

Given the emphasis in our suppressivity model on repeat unit size, but also notable outliers such as family 2 in Figure 8b, it is also possible that effective repeat unit sizes dictated by secondary structure play an important role. Suppressivities in Petite family 2, which contain tandemly repeated inverted dimers, deviate most from the theoretical curve. Inverted sequences like those in family 2 would also be expected to form the hairpin structure hypothesized in the generative model of the mixed repeats. If this hairpin persists, it would consume directly repeated regions that are preferred in crossover events. The result would be a reduction in the density of repeated regions accessible to excision events, which upon eventual fragmentation through excisions would produce a larger effective repeat unit. This may in part explain the deviation of Petite family 2 from the theoretical suppressivity, as well as the inverted repeat rule itself that was proposed in (de Zamaroczy et al., 1981). Overall, more data is likely required to aggregate these apparent exceptions or outliers into a more encompassing model of suppressivity, but the present study provides a foundation of modern techniques and lessons to build upon in this goal.

Finally, it’s worthwhile to comment on the tolerance of the Petite mutation in populations and why their mtDNA might have evolved to produce a structure susceptible to a wide array of destructive excision events. The confirmation of the rapid recombination hypothesis in the generation of mixed structures in this study demonstrates the high frequency of recombination of mtDNA in yeast. The destructiveness of the Petite mutation in the background of this frequent recombination illustrates the danger of sequence specific illegitimate recombination in this system. While the evolutionary origin of the repetitiveness of the mitochondrial genome of yeast has been explored and the hypothesis put forth that the repetitive origins and surrogate origin sequences confer a replication or transcriptional advantage (de Zamaroczy & Bernardi, 1986; Bernardi, 2005), it is clear that this structure is accompanied by a trajectory of continued fragmentation without external forces opposing this fate. The persistence of this repetitive structure in *S. cerevisiae* strongly suggests that both the genome repetition and rapid recombination that follows from it was an evolutionary product of strong intracellular selection in conjunction with strong population level selection for respiring yeasts with intact genomes. Thus, both genome multiplicity and these opposing levels of selection were necessary to evolve this mtDNA structure so susceptible to self-destruction, but also allow for an especially high recombination rate from which cells could benefit through the genetic complementation of errors and perhaps other uses such as the destruction of invasive DNA. As such, mtDNA dynamics and the Petite mutation in yeast is a wonderful example of how multi-level selection has shaped evolutionary trajectories of genomes.

## Methods

### Details of experiments

#### Yeast strains and their construction for suppressivity testing

The Grande tester strain used in mating with Petites was the baker’s yeast strain yCO362 a W303 leu2-3,112 can1-100 ura3-1 ade2-1 his3-11,15, which was a gift from the Boris Shraiman lab at UCSB. To construct the Grande progenitor of Petite strains, we restored URA3 function in SY2081 α W303 leu2-3,112 can1-100 ura3-1 ade2-1 his3-11,15 trp1-1 which was a gift from the Grant Brown lab at the University of Toronto. To this end, we grew an E. coli strain harbouring pFA6a-URA3, which was a gift from Jon Houseley & David Tollervey (Addgene # 61924). Plasmids were extracted and the URA3 fragment PCR amplified with primers that share 20nt of short flanking homology with the reference yeast mitochondrial genome following the standard short flanking homology targeted recombination method (Petracek & Longtine, 2002). Expected PCR fragment sizes were confirmed on a gel and then transfected into SY2081 using the high efficiency LiOAC yeast transformation protocol (Brown et al., 2015). Integration at the expected location was confirmed through PCR of flanking regions overlapping each breakpoint, and Sanger sequencing. To ensure integration was exclusive to our target location we then performed tetrad analysis on the transformed SY2081 (named 10T3 hereafter) x yCO362 and observed 2:2 segregation as expected for a single integration site.

#### Media and growth conditions

Both Grande and Petite colonies were cultured in YPAD medium (1% yeast extract, 2% bacto-peptone, 2% glucose, 0.072% adenine hemisulfate). Petite colonies were detected under growth in YPADG medium (1% yeast extract, 2% bacto-peptone, 0.1% glucose, 3% glycerol, 0.072% adenine hemisulfate). With a reduced glucose content, Petite colonies appear smaller and more translucent than their Grande counterparts in this media and the differences were discernible beyond 4 days at 30°C (Dimitrov et al., 2009). YPG medium (1% yeast extract, 2% bacto-peptone, 3% glycerol) was used in culturing a subset of Grande colonies, and verifying the respiratory deficiency of identified Petites. To measure suppressivity, we used SC-ura-trp (DG carbon source) media (0.67% bacto yeast nitrogen base w/o amino acids, 0.1% glucose, 3% glycerol, 0.2% dropout powder lacking uracil and tryptophan) which selects for zygotes due to strain auxotrophies. Liquid cultures were grown at 30°C in a linear shaking water bath, while solid media growth took place in a forced air incubator also at 30°C.

#### Isolation of spontaneous petite colonies

Liquid culture of strain 10T3 inoculated in YPAD media was washed in dH2O and plated on YPADG media. Respiratory deficiency of these Petite colonies was confirmed through replica plating on to YPG, as well as patching onto separate YPG plates. Following confirmation of these colonies being Petites, 9 different colonies were streaked onto YPAD agar as a first passage. Three colonies were randomly selected from this first passage plate and streaked again onto YPAD agar, constituting a second passage. Three colonies from each second passage plate were cultured, stored as frozen stocks, had their suppressivities measured, and a subset were sequenced.

#### Yeast suppressivity assay

Cultures of Grande (yCO362) and Petite strains (10T3 derived) were grown overnight in YPAD liquid media. Cultures were diluted to 0.1OD and grown for three hours. Equal volumes of each culture were mixed and incubated at room temperature for 20 hours to allow for mating. A small aliquot of this mating mixture was observed under a hemocytometer to calculate appropriate dilutions for plating. The mating mixture was diluted and washed in dH20, then plated onto SC-ura-trp (DG carbon source). After 5 days at 30°C Petri dishes were scanned, and the fraction of small to total colonies on these plates was recorded as the suppressivity of the strain, calculated based on an average of 250 colonies labeled per strain. The average standard deviation in suppressivity across strains related by the same first passage progenitor was ~ 5%.

#### Nanopore sequencing

Three or more progeny from 9 separate spontaneous Petites derived from strain 10T3, and 10 Grande colonies under YPD (6) /YPG (4) liquid culture growth of the same strain were sequenced on a Oxford Nanopore MinION Mk1B (Figure 1). Whole genomic DNA from these 48 colonies following culturing was extracted using a modified enzymatic Hoffman-Winston DNA extraction protocol as described in (Boeke et al., 1985). DNA was then purified with a Qiagen 20/G Genomic-tip, and barcoded with the Oxford Nanopore EXP-NBD104 and EXP_NBD114 barcoding kits in conjunction with the SQK-LSK109 ligation sequencing kit with long fragment Agencourt AMPureXP purification, following manufacturer’s instructions. Twenty-four barcoded samples were pooled at a time in two FLO-MIN106 flow cells with R9.4.1 chemistry. Sequencing generated a total of 8.5Gbases of reads across both flow cells within 24 hours, with a mean read length of ~6kbp and maximum length of 120kbp. 429 Mbases of reads were mapped to mitochondrial DNA, resulting in an average coverage per primary structure of ~700 across all Petite samples sequenced.

### Details of analysis

#### Read basecalling, alignment, and filtering

Raw Nanopore reads were basecalled and demultiplexed with the ONT Guppy package 3.1.5-1. Reads that passed default quality score filtering (>9 qscore) were aligned using Minimap2 with default parameters (Li, H., 2018) to the *S. cerevisiae* reference release R64-2-1_20150113 (Engel et al., 2014) available from yeastgenome.org. Following initial alignment, unmapped regions in reads were recursively aligned to the reference sequence to combat the Z-drop heuristic, which exists to remove spurious alignment artifacts introduced due to the expectation of collinearity between alignment anchors. This is a particularly insidious heuristic when mapping Petite structures due to large numbers of repeats within reads. See Figure S11, where we provide an example of the effect of this Z-drop heuristic in resolving Petite structures, and how in a small subset of reads the pseudo-global alignment produced in our recursive mapping improves structure resolution. Alignments in reads were kept with PHRED scaled mapping quality scores > 20, and alignment lengths >300bp due to the high degree of homology between mt replication origins which makes alignment of these small regions of context difficult (Zamaroczy et al., 1981).

#### Mitochondrial DNA alignment breakpoint detection

Alignment breakpoints are defined in our pipeline to be deviations of more than 30bp between the read and reference coordinates in adjacent alignments within one read. They are also identified by strand changes across adjacent alignments regardless of the separation in reference/read coordinates. As such, these breakpoint signals encompass large insertions, and deletions, and inversions within reads, and are delineated by alignment termination from the mapping program Minimap2 (with default parameters) which are stored and processed in a structural detection pipeline written in Python.

#### Removal of prevalent inverted duplication artifacts

Before breakpoint signals can be clustered, inverted duplication artifacts, which are prevalent in our particular sequencing chemistry and are apparently affected by growth media (Figure S1a), must be filtered out from real inverted duplication breakpoint signals. These artifacts appear to be due to complementary strands being pulled in succession through the pore, resulting in an unfolding of a double stranded DNA molecule into a molecule of double the length with a characteristic inversion. This may be due to either physical tethering, or lingering of the separated complementary strand near the pore opening. In any case, conveniently this results in a nearly centered inversion within such reads (Figure S1b), that is sometimes skewed towards the end of the read because of increased translocation speed and self-interaction of the strands in a ratcheting mechanism as described in (Spealman et al., 2020). This increased speed also results in reduced read quality in the latter half as reported in (Spealman et al., 2020) and Figure S5a, which means that distances between reference positions of alignments at inversions have the distribution in Figure S5b. Given that 90% of artifacts result in adjacent alignment edges with distances less than 1000bp we take this to be one of our criteria for an inversion artifact.

Inverted duplication artifacts are filtered from real inverted duplications at two levels before clustering: (1) Breakpoint signals where within read positioning of the inverted breakpoint is > 1% likely to be derived from the purely artifact distribution in Figure S1b is recorded in a list. (2) If breakpoints that follow criteria (1) are derived from reads with only a single inverted duplication signal, then they are considered artifacts.

#### Breakpoint clustering

Inverted/non-inverted breakpoints are clustered separately using the DBSCAN algorithm (Ester et al., 1996). This algorithm requires two parameters: ∈, which defines the neighbourhood of a breakpoint as a radius in base pairs, and *min Pts*, which is the number points required within *ϵ* of a breakpoint to be considered a core point, or dense region. Breakpoints within dense regions are then connected together iteratively to produce a density-connected cluster.

For both inverted and non-inverted breakpoints, we set *min Pts* = 3, to capture even the smallest clusters which will undergo further filtering described later. For non-inverted breakpoints, *ϵ* = 1*kbp*, which is a lenient choice given that the most catastrophic deviations in expected breakpoint positions due to sequencing error in inverted duplication artifacts are largely under 1kbp. For inverted breakpoints post artifact-filtering described above, an optimal *ϵ* is computed by sorting nearest neighbour distances across all inverted breakpoints, and computing the Euclidean distance to the nearest neighbour that results in the largest curvature in a plot of nearest neighbour distance vs breakpoint number. This is the so-called “elbow” method. Then *ϵ* = *min*(*optimal*(*ϵ*), 1*kbp*, is taken to be the nearest-neighbour radius for inverted breakpoints. This extra step for inverted breakpoints is required solely because of the possibility of low frequency artifact noise that is not present for non-inverted breakpoints.

#### Breakpoint filtering based on read support and a majority voting scheme

Following clustering with the above parameters, breakpoint clusters are then required to have a minimum of 3 separate reads supporting them to be considered real. Furthermore, in reads that contain a detected inverted duplication artifact, we use the duplicated signals on either side of this artifact to our advantage in a majority voting scheme: For breakpoints belonging to cluster *j*, we count the number of times a breakpoint belonging to *j* is recapitulated on either side of an inverted duplication artifact, *P_j_* and the number of times it isn’t, *N_j_*. If *P_j_* > *N_j_* we consider this breakpoint to be real. In the case that *P_j_* = *N_j_* = 0, meaning that the breakpoint doesn’t exist in reads that contain inverted duplication artifacts, we perform a similar majority vote on the basis of a breakpoint being repeated or not within reads. The notion here is that real breakpoints should be repeated within reads because of the expected concatemer structure of Petites, while spurious breakpoints that are low frequency are less likely to be. To this end we compute *NR_j_* and *PR_j_*, which are the number of times a breakpoint from cluster *j* is present in a read with repeats but is not repeated itself, and the number of times a breakpoint assigned to cluster *j* is repeated, respectively. Similar to the above, if *PR_j_* > *NR_j_* we consider this cluster to be real.

See Table S1 for a summary of all parameters in our pipeline described in this section and the three preceding it. See Figure S12 for the effect that changing these parameters has on breakpoint counts. See Figure S10 for the effect that this majority voting scheme has on removing spurious breakpoints.

#### Breakpoint labeling and read encoding schemes

Breakpoints are labeled based on the segmentation provided by a Python implementation of DBSCAN (an integer), in addition to two other features: (1) the strand in this transition (+/-), and (2) the transition from a low (L) to high (H) or vice-versa reference base position. This means that read with a repeated excised repeat unit with the same orientation and in the + strand takes the form: [2LH, 2LH, 2LH,..,2LH] in the number transition encoding, and [2++, 2++, 2++,…,2++] in the orientation encoding scheme. In addition to these two schemes, the mapped ends of the reads in reference locations are also stored and used later in assigning reads to inferred structures. Both encoding schemes are necessary when considering complex structures such as the “mixed” repeat structures described in results where only two breakpoints exist, but permutations in their orientation produce four unique alignments which would be missed with a simple numerical labeling scheme.

#### Reconstructing mtDNA repeat structures from reads

For long reads containing small repeated structures with numerous breakpoints passing filtering criteria, repeats are detected by directly computing the longest common prefix within a read in both read encoding schemes described above. We require two periods to be present for this type of repeat detection method to produce a candidate structure, and that the detected repeat is a tandem repeat, meaning there are no intervening breakpoints between periods. This is how all low frequency alternate structures are detected within samples, and for some samples primary structures when small enough to be repeated within single reads.

Large primary structures that are too large to be fully repeated twice within reads are inferred through the construction of a breakpoint transition matrix, *T_ij_*, that stores the number of transitions between breakpoint *i* to breakpoint *j* across all reads within a sample. If two entries in this matrix share a breakpoint, and if the fractional difference in their average counts to individual counts are < 0.34, these breakpoint transitions are merged into a list together. This is a Binomial merging criterion for being within one standard deviation of the average counts assuming that both breakpoint transitions sharing a breakpoint come from the same structure. Following recursive merging of transitions and then lists of transitions with the same criteria, if in each list the number of breakpoints equals the number of unique transitions (meaning this structure is a true repeat), then the structure is recorded alongside its average count and its span on the reference. Structures inferred from this method with the largest span on the reference are considered to be primary repeats, and if small enough are recapitulated by the direct repeat detection scheme described above.

#### Mixed structure alignment frequency calculations

Consider an alignment of length *L*_1_, where we are interested in computing its frequency in these mixed samples that contain four alignments with lengths {*L*_1_, *L*_2_, *L*_3_, *L*_4_}, accounting for read sampling bias. Given a read of length *x* > *L*_1_, assuming sampling of a random pattern of all four alignments (or equivalently, random sampling of a periodic pattern including all four alignments), the probability this read isn’t truncating an alignment with length *L*_1_ is 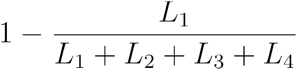.

The estimated probability that we see this read of length *x* > *L*_1_ is:

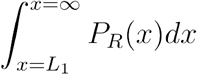

Where *P_R_*(*x*) is an exponential fit to the empirical read length distribution (Figure S13), with following form, and is the read length probability distribution:

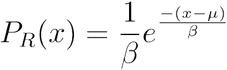

Where *β* and *μ* are the scale and location parameters determined from the fit.

The joint probability that we have a read long enough to see an alignment, and that it does not abruptly truncate the alignment at its ends is the product of these two terms:

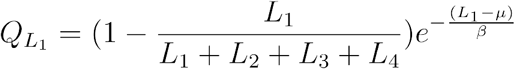

Alignment frequencies can then be computed by normalizing raw observed counts *N_i_* by *Q_L_i__*, and computing their relative frequency, *ν_i_*:

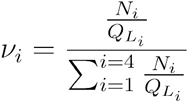

#### Primary/alternate structure frequency calculations

Unlike mixed samples where breakpoint identity does not uniquely define alignments (permutations of two breakpoints result in 4 alignments), in most samples we can simply count breakpoints, assuming that repeat units ~ unique breakpoint counts in a particular structure, and perform a similar normalization to account for sampling bias.

To do this, each read is assigned to one class of potential structures based on the maximal overlap between breakpoint labels in known structures and the read, conditioned on containing content only within the known structure. Because we are counting breakpoints, we no longer have to worry about truncation of a whole alignment, and can consider reads smaller than the expected alignment length.

Consider a repeat unit with period *L_k_* in bp, and a breakpoint count *J_k_* that is unique to this structure. The probability that we see this structure given the read distribution in this sample is:

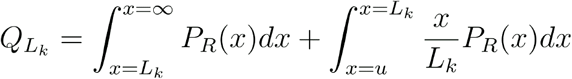

The first term is the product of the probability (=1) that we see this breakpoint for read lengths *x* > *L_k_* and the probability that we have such reads (*P_R_*(*x*) is the read-length distribution fit as described above). The second term is the product of the probability we see this breakpoint for reads 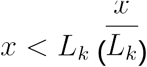 and the probability that we see reads with these lengths.

The relative frequency can then be calculated by normalizing counts *J_k_* by *Q_L_k__* and computing relative frequency across all structures as in the previous section.

#### Type I/II/III repeat unit classification

Type I repeat units are those excised from the interior of a primary Petite alignment. Type II repeat units contain alignments that span the primary alignment breakpoint when it is present in a concatemer form. Type III repeat units share one edge with the primary alignment, and have another edge internal to the primary alignment. Because each edge in a reconstructed structure is truly a cluster of edge signals, for an alignment edge to be considered “shared” the means of alignment edge location distributions must be equal to within one standard deviation of each other. Thus, according to this criteria, Type I repeats have no shared edges, Type II repeat units have two such shared edges and multiple alignments, and Type III repeat units have one shared edge and one alignment.

## Acknowledgements

We thank Dr. Grant Brown at the University of Toronto for his thorough reading of the manuscript and numerous members of the Goyal lab for their thoughtful comments and discussions. We also thank Dr. Boris Shraiman and Dr. Grant Brown for yeast strains. We recieved funding from the Natural Sciences and Engineering Research Council of Canada (NSERC) Discovery Grant RGPIN-2015-0, the Simons Foundation Grant 326844 in the Mathematical Modeling of Living Systems, and funding for equipment from the Canadian Foundation for Innovation (CFI) Grant 32708.

## Notes

### Competing Interest Statement

The authors have declared no competing interest.

### Summary of Updates

Formatting, simplification of language, and supplemental files updated.

## References

Baldacci, G., & Bernardi, G. (1982). Replication origins are associated with transcription initiation sequences in the mitochondrial genome of yeast. The EMBO Journal, 1(8), 987–994. https://doi.org/10.1002/j.1460-2075.1982.tb01282.x

Bernardi, G., Prunell, A., Kopecka, H. (1975). An analysis of the mitochondrial genome of yeast with restriction enzymes. In: Puiseux-Dao, S. (Ed.), Molecular Biology of Nucleocytoplasmic Relationships. Elsevier, Amsterdam, The Netherlands, pp. 85–90.

Bernardi, G., Prunell, A., Fonty, G., Kopecka, H., Strauss, F. (1976). The mitochondrial genome of yeast: organization, evolution, and the petite mutation. In: Saccone, C., Kroon, A.M. (Eds.), Proceedings of the 10th International Bari Conference on the Genetic Function of Mitochondrial DNA. Elsevier North-Holland, Amsterdam, The Netherlands, pp. 185–198.

Bernardi, G., Bernardi, G. (1980). Repeated sequences in the mitochondrial genome of yeast. FEBS Lett. 115, 159–162.

Bernardi, G. (2005). Lessons from a small, dispensable genome: The mitochondrial genome of yeast. Gene, 354(1-2 SPEC. ISS.), 189–200. https://doi.org/10.1016/j.gene.2005.03.024

Blanc, H., & Dujon, B. (1980). Replicator regions of the yeast mitochondrial DNA responsible for suppressiveness. Proceedings of the National Academy of Sciences of the United States of America, 77(7 II), 3942–3946.

Boeke JD, Garfinkel DJ, Styles CA, Fink GR. (1985). Ty elements transpose through an RNA intermediate. Cell. 1985 Mar;40(3):491–500. doi: 10.1016/0092-8674(85)90197-7. PMID: 2982495.

Bos JL, Heyting C, Van der Horst G, Borst P. (1980). The organization of repeating units in mitochondrial DNA from yeast petite mutants. Curr Genet. 1980 Apr;1(3):233–9. doi: 10.1007/BF00390949. PMID: 24189664.

Bradshaw, E., Yoshida, M., Ling, F. (2017). G3: GENES, GENOMES, GENETICS September 1, 2017 vol. 7 no. 9 3083–3090; https://doi.org/10.1534/g3.117.043851

Brewer BJ, Payen C, Raghuraman MK, Dunham MJ (2011) Origin-Dependent Inverted-Repeat Amplification: A Replication-Based Model for Generating Palindromic Amplicons. PLOS Genetics 7(3): e1002016. https://doi.org/10.1371/journal.pgen.1002016

Brewer BJ, Payen C, Di Rienzi SC, Higgins MM, Ong G, et al. (2015) Origin-Dependent Inverted-Repeat Amplification: Tests of a Model for Inverted DNA Amplification. PLOS Genetics 11(12): e1005699. https://doi.org/10.1371/journal.pgen.1005699

Brown GW., Dunham MJ., Gartenberg MR. (2015). Methods in Yeast Genetics and Genomics: High-efficiency Yeast Transformation. Cold Spring Harbor Laboratory Press.

Chan, D. C. (2006) Mitochondria: Dynamic Organelles in Disease, Aging, and Development. Cell 125, 1241–1252.

de Zamaroczy, M., Baldacci, G., Bernardi, G. (1979). Putative origins of replication in the mitochondrial genome of yeast. FEBS Letters 108, 429–432.

de Zamaroczy, M., Marotta, R., Faugeron-fonty, G., Goursot, R., Mangin, M., Baldacd, G., & Bernardi, G. (1981). The origins of replication of the yeast mitochondrial genome and the phenomenon of suppressivity. Nature, 292(July), 0–3.

de Zamaroczy, M., Faugeron-Fonty, G., & Bernardi, G. (1983). Excision sequences in the mitochondrial genome of yeast. Gene, 21(3), 193–202. https://doi.org/10.1016/0378-1119(83)90002-1

de Zamaroczy, M., & Bernardi, G. (1986). The GC clusters of the mitochondrial genome of yeast and their evolutionary origin. Gene, 41(1), 1–22. https://doi.org/10.1016/0378-1119(86)90262-3

Dimitrov LN, Brem RB, Kruglyak L, Gottschling DE. Polymorphisms in multiple genes contribute to the spontaneous mitochondrial genome instability of Saccharomyces cerevisiae S288C strains. (2009). Genetics. 2009 Sep;183(1):365–83. doi: 10.1534/genetics.109.104497. Epub 2009 Jul 6. PMID: 19581448; PMCID: PMC2746160.

Engel, S. R., Dietrich, F. S., Fisk, D. G., Binkley, G., Balakrishnan, R., Costanzo, M. C., Dwight, S. S., Hitz, B. C., Karra, K., Nash, R. S., Weng, S., Wong, E. D., Lloyd, P., Skrzypek, M. S., Miyasato, S. R., Simison, M., & Cherry, J. M. (2014). The reference genome sequence of Saccharomyces cerevisiae: then and now. G3 (Bethesda, Md.), 4(3), 389–398. https://doi.org/10.1534/g3.113.008995

Ephrussi, B., (1949). Action de l’acriflavine sur les levures. Unite’s biologiques doue’es de continuite’ge’ne’tique VIII. Colloques internatio-naux du CNRS-Publications du CNRS, Paris, France.

Ephrussi, B., Hottinguer, H., Tavlitzki, J. (1949). Action de l’acriflovine sur les levures. II-Etude genetique de mutant “petite colonie.” Ann. Inst. Pasteur (Paris), 77, pp. 419–450

Ephrussi, B. (1953). Nucleo-cytoplasmic relations in micro-organisms— their bearing on cell heredity and differentiation. Oxford at the Clarendon Press, Oxford, UK.

Ester, Martin; Kriegel, Hans-Peter; Sander, Jörg; Xu, Xiaowei (1996). Simoudis, Evangelos; Han, Jiawei; Fayyad, Usama M. (eds.). A density-based algorithm for discovering clusters in large spatial databases with noise. Proceedings of the Second International Conference on Knowledge Discovery and Data Mining (KDD-96). AAAI Press. pp. 226–231. CiteSeerX 10.1.1.121.9220. ISBN 1-57735-004-9.

Faugeron-Fonty, G., Culard, F., Baldacci, G., Goursot, R., Prunell, A., & Bernardi, G. (1979). The mitochondrial genome of wild-type yeast cells. VIII. The spontaneous cytoplasmic “petite” mutation. Journal of Molecular Biology, 134(3), 493–537. https://doi.org/10.1016/0022-2836(79)90365-6

Faugeron-Fonty G, Mangin M, Huyard A, Bernardi G. (1983). The mitochondrial genomes of spontaneous orir petite mutants of yeast have rearranged repeat units organized as inverted tandem dimers. Gene. 1983 Sep;24(1):61–71. doi: 10.1016/0378-1119(83)90131-2. PMID: 6354845.

Fayet, G., Jansson, M., Sternberg, D., Moslemi, A., Fardeau, M., Oldfors, A., … Lombe, A. (2002). Ageing muscle: clonal expansions of mitochondrial DNA point mutations and deletions cause focal impairment of mitochondrial function. Neuromuscul Discord, 12, 484–493.

Foury, F., Roganti, T., Lecrenier, N., Purnelle, B. (1998). The complete sequence of the mitochondrial genome of Saccharomyces cerevisiae. FEBS Lett. 440, 325–331.

Fritsch, E. S., Chabbert, C. D., Klaus, B., & Steinmetz, L. M. (2014). A genome-wide map of mitochondrial DNA recombination in yeast. Genetics, 198(2), 755–771. https://doi.org/10.1534/genetics.114.166637

Gaillard, C., Strauss, F. & Bernardi, G. (1980). Excision sequences in the mitochondrial genome of yeast. Nature 283, 218–220. https://doi.org/10.1038/283218a0

Goursot, R., de Zamaroczy, M., Baldacci, G., Bernardi, G. (1980). Super-suppressive petite mutants in yeast. Curr. Genet. 1, 173–176.

Goursot, R., Mangin, M., & Bernardi, G. (1982). Surrogate origins of replication in the mitochondrial genomes of ori-zero petite mutants of yeast. The EMBO Journal, 1(6), 705–711. https://doi.org/10.1002/j.1460-2075.1982.tb01234.x

Heyting C, Talen JL, Weijers PJ, Borst P. (1979). Fine structure of the 21S ribosomal RNA region on yeast mitochondrial DNA. II. The organization of sequences in petite mitochondrial DNAs carrying genetic markers from the 21S region. Mol Gen Genet. 1979 Jan 11;168(3):251–77. doi: 10.1007/BF00271497. PMID: 374988.

Holt, Ian J. (2019). The mitochondrial R-loop, Nucleic Acids Research, Volume 47, Issue 11, 20 June 2019, Pages 5480–5489, https://doi.org/10.1093/nar/gkz277

Kozik, A., Rowan, B. A., Lavelle, D., Berke, L., Eric Schranz, M., Michelmore, R. W., & Christensen, A. C. (2019). The alternative reality of plant mitochondrial DNA: One ring does not rule them all. PLoS Genetics, 15(8), 1–30.https://doi.org/10.1371/journal.pgen.1008373

Lewin, A., Morimoto, R., Rabinowitz, M., & Fukuhara, H. (1978). Restriction enzyme analysis of mitochondrial DNAs of petite mutants of yeast: Classification of petites, and deletion mapping of mitochondrial genes. MGG Molecular & General Genetics, 163(3), 257–275. https://doi.org/10.1007/BF00271955

Lewin, A. S., Morimoto, R., and Rabinowitz, M. (1979). Stable heterogeneity of mitochondrial DNA in grande and petite strains of S. cerevisiae. Plasmid 2, 474–484.

Li, H. (2018). Minimap2: Pairwise alignment for nucleotide sequences. Bioinformatics, 34(18), 3094–3100. https://doi.org/10.1093/bioinformatics/bty191

Ling, F., & Shibata, T. (2002). Recombination-dependent mtDNA partitioning: In vivo role of Mhr1p to promote pairing of homologous DNA. EMBO Journal, 21(17), 4730–4740. https://doi.org/10.1093/emboj/cdf466

Ling, F., & Shibata, T. (2004). Mhr1p-dependent Concatemeric Mitochondrial DNA Formation for Generating Yeast Mitochondrial Homoplasmic Cells □, 15(January), 310–322. https://doi.org/10.1091/mbc.E03

Ling, F., Hori, A., & Shibata, T. (2007). DNA Recombination-Initiation Plays a Role in the Extremely Biased Inheritance of Yeast [rho-] Mitochondrial DNA That Contains the Replication Origin ori5. Molecular and Cellular Biology, 27(3), 1133–1145. https://doi.org/10.1128/mcb.00770-06

Ling, F., Bradshaw, E., Yoshida, M. (2019). Prevention of mitochondrial genomic instability in yeast by the mitochondrial recombinase Mhr1. Sci Rep 9, 5433. https://doi.org/10.1038/s41598-019-41699-9

Locker, J., Rabinowitz, M., & Getz, G. S. (1974). Tandem inverted repeats in mitochondrial DNA of petite mutants of Saccharomyces cerevisiae. Proceedings of the National Academy of Sciences of the United States of America, 71(4), 1366–1370. https://doi.org/10.1073/pnas.71.4.1366

Locker, J., Lewin, A., & Rabinowitz, M. (1979). The structure and organization of mitochondrial DNA from petite yeast. Plasmid, 2(2), 155–181. https://doi.org/10.1016/0147-619X(79)90036-2

MacAlpine, D. M., Kolesar, J., Okamoto, K., Butow, R. A., & Perlman, P. S. (2001). Replication and preferential inheritance of hypersuppressive petite mitochondrial DNA. The EMBO journal, 20(7), 1807–1817. https://doi.org/10.1093/emboj/20.7.1807

Mangin, M., Faugeron-Fonty, G., & Bernardi, G. (1983). The orir to ori+ mutation in spontaneous yeast petites is accompanied by a drastic change in mitochondrial genome replication. Gene, 24(1), 73–81. https://doi.org/10.1016/0378-1119(83)90132-4

Marotta, R., Colin, Y., Goursot, R., & Bernardi, G. (1982). A region of extreme instability in the mitochondrial genome of yeast. The EMBO Journal, 1(5), 529–534. https://doi.org/10.1002/j.1460-2075.1982.tb01204.x

Osman, C., Noriega, T. R., Okreglak, V., Fung, J. C., & Walter, P. (2015). Integrity of the yeast mitochondrial genome, but not its distribution and inheritance, relies on mitochondrial fission and fusion. Proceedings of the National Academy of Sciences of the United States of America, 112(9), E947–956. https://doi.org/10.1073/pnas.1501737112

Payne, B. & Chinnery, P. (2015) Mitochondrial dysfunction in aging: Much progress but many unresolved questions. Biochimica et Biophysica Acta 1847, 1347–1353.

Petracek ME, Longtine MS. (2002). PCR-based engineering of yeast genome. Meth Enzymol 350: 445–469.

Rayko, E., Goursot, R., Cherif-Zahar, B., Melis, R., & Bernardi, G. (1988). Regions flanking ori sequences affect the replication efficiency of the mitochondrial genome of ori+ petite mutants from yeast. Gene, 63(2), 213–226. https://doi.org/10.1016/0378-1119(88)90526-4

Spealman, P., Burrell, J., Gresham, D. (2020) Inverted duplicate DNA sequences increase translocation rates through sequencing nanopores resulting in reduced base calling accuracy, Nucleic Acids Research, Volume 48, Issue 9, 21 May 2020, Pages 4940–4945, https://doi.org/10.1093/nar/gkaa206

Veatch JR, McMurray MA, Nelson ZW, Gottschling DE. (2009). Mitochondrial dysfunction leads to nuclear genome instability via an iron-sulfur cluster defect. Cell. 2009 Jun 26;137(7):1247–58. doi: 10.1016/j.cell.2009.04.014. PMID: 19563757; PMCID: PMC2759275.

