## Supplementary figures and tables for "Contingency and selection in mitochondrial genome dynamics"

### Supplementary Information: Figures

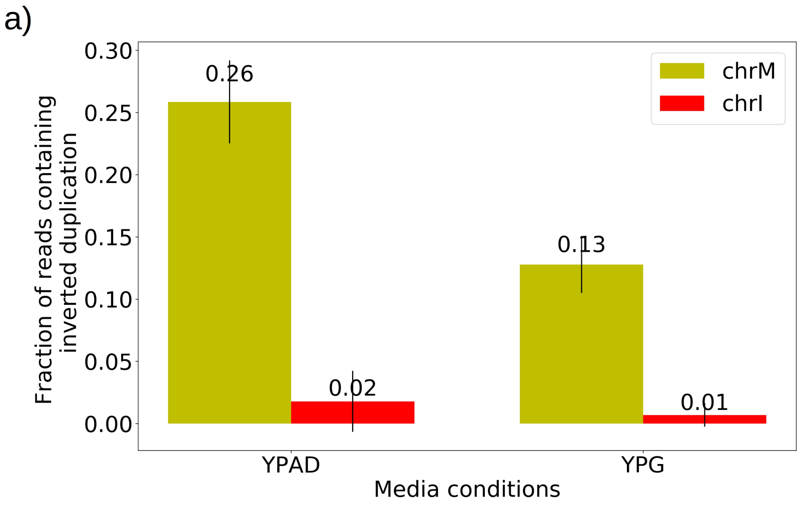


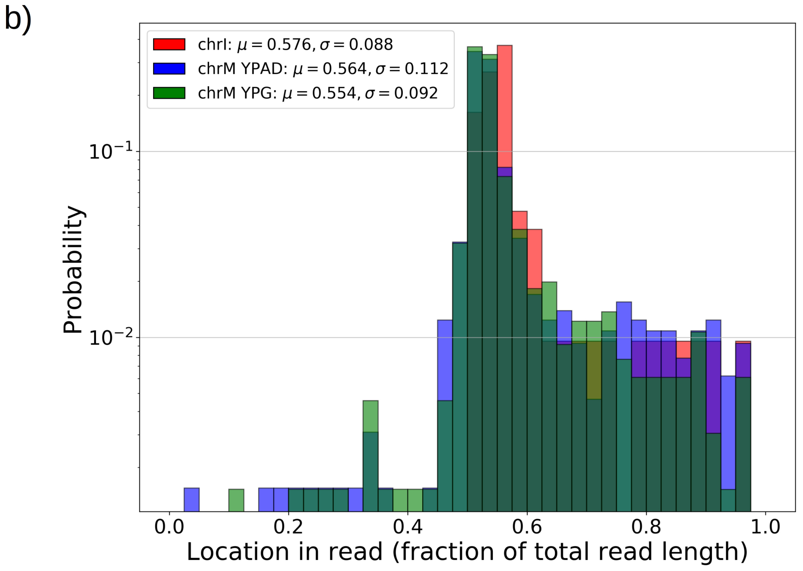


**Figure S1: Inverted duplication artifacts are prevalent in Nanopore sequencing of mitochondrial DNA but exhibit patterns that enable their detection**

a) The fraction of reads containing repeated inverted annotated genome features were plotted above for all Grande colonies grown in YPD and YPG for both nuclear DNA (nDNA) and mtDNA. Inverted duplication artifacts were enriched in reads mapped to mitochondrial DNA. While it has been suggested that these artifacts are caused by either tethered complementary strands being brought through the pore in series, or lingering complementary strands near the pore opening, it is unclear why there is such a difference between nDNA and mtDNA reads. Base composition does differ significantly between the two genomes, with only 18% GC content in mtDNA, compared to 38% in nDNA. The abundance of inverted duplication artifacts also appears to be affected by growth conditions, potentially due to differences in mtDNA conformation under respiration and fermentation conditions, but this remains unexplored. This effect was also seen in other Nanopore experiments performed by another lab with the same flow cell and sequencing chemistry described in methods (data not shown). b) Here we are plotting the probability distribution of the read locations of singleton inverted duplication artifacts in mtDNA across all Grande samples for both YPG and YPD conditions, as well as pooled reads from chromosome I (chrI) across both conditions. Mean and standard deviations of each set of samples are denoted in the legend. Artifact inverted duplications are generally concentrated towards the center of the read, but biased slightly towards the second half. This is consistent with the self-interaction ratcheting mechanism described by (Spealman et al., 2020) in sequencing real inverted duplications, where self interaction increases translocation speed in the second half of the read. Increased translocation speed results in skipped bases, effectively shortening the second half of the read which results in this bias to the right in fractional length. Inverted duplications detected are filtered to those residing within the 1% tails of this distribution.

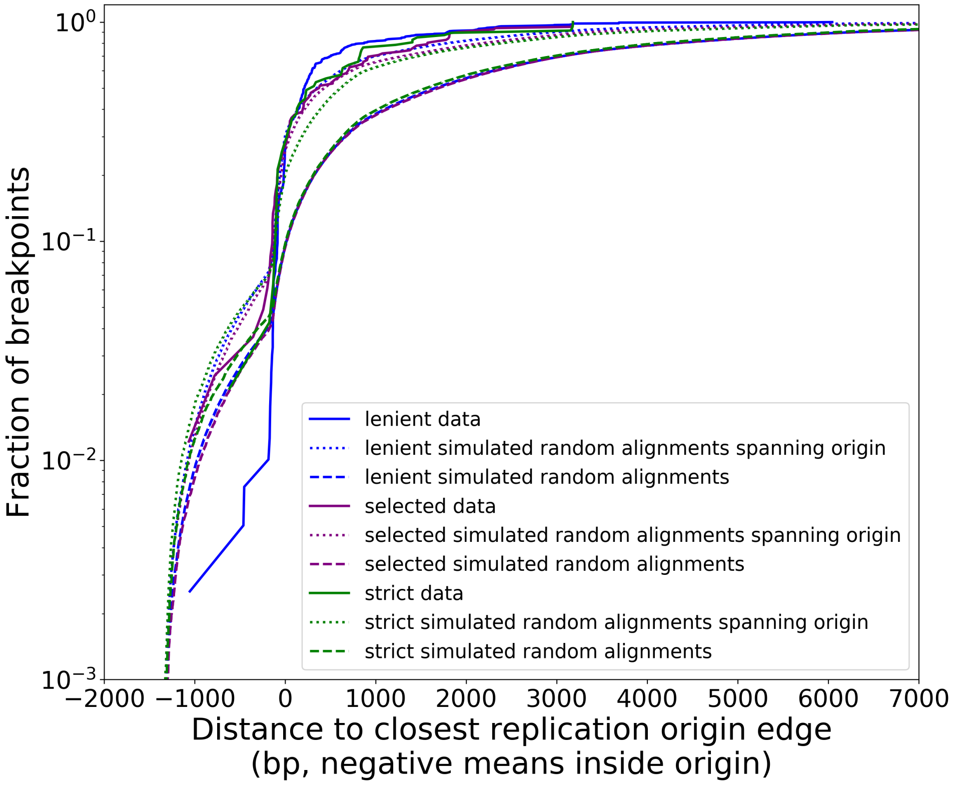


**Figure S2: Distribution of minimum distances between alignment edges and origin edges is robust to different structural detection pipeline parameter regimes**

Real data (solid lines in this plot) follow all filtering steps in our structural pipeline, including inverted duplication artifact filtering, and majority voting (except “lenient” parameter set). The different data curves are a result of the aforementioned filtering steps in the pipeline, but with different parameters (Table S2). Small dotted lines represent simulations of uniform random alignments spanning origins, with length distributions of alignments from each set of data curves. Long dashed lines represent the same type of simulation with no requirement for alignments to span origins. Both the green and purple data curves reside close to the “strong origin selection” models or the small dotted lines. The blue parameter regime, which we would expect to cluster more noise because we are being more lenient with filtering thresholds, differs at least to a larger degree than green/purple. Overall, however, all three parameter regimes perform similarly, suggesting that the shape of these distributions and the claims we are making here are robust at least to changes in parameter values in our structure detection pipeline. In another sense, this suggests that minimap2 is already neglecting most base-level changes very well and only considering severe deviations in expected collinearity to be the end of alignments that form breakpoints.

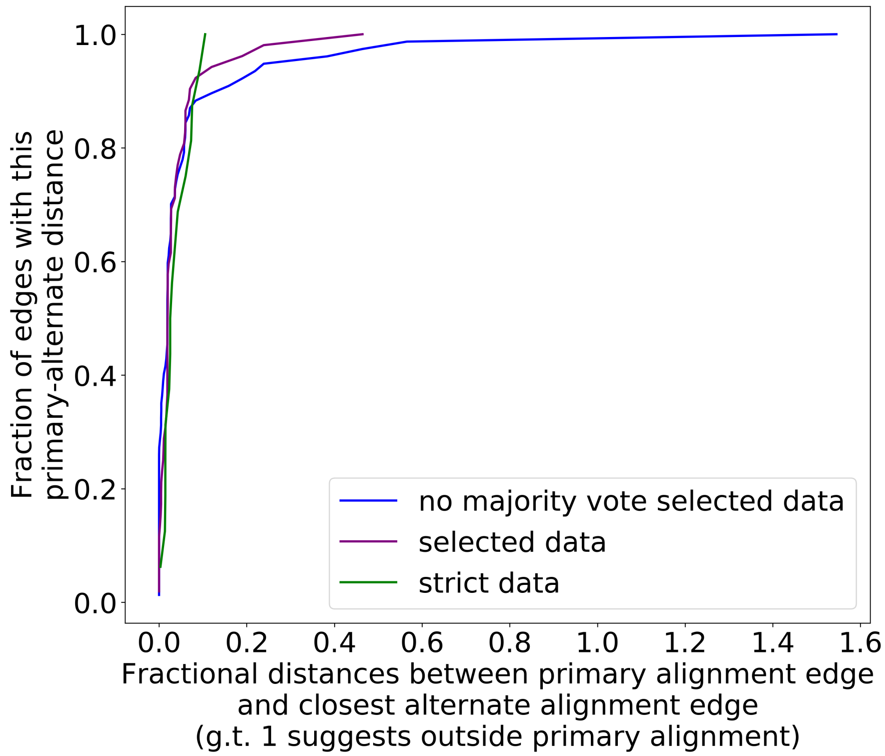


**Figure S3: The preference for primary structure excision sites is robust to different filtering parameter regimes**

Here we are plotting the cumulative distribution of fractional distances between the primary alignment edge and closest alternate alignment edge for three different parameter regimes (Table S2). The blue curve, which represents our “lenient” regime in this case, is now simply the purple curve (parameters in Table S1, regimes in Table S2) without majority voting. This change is necessary compared to Figure S2 because we now must infer structure through repeat detection, which requires more strict parameters to begin with. Without the majority voting it is clear that alternate alignments begin to creep outside primary alignments due to noise, but as a whole, all three parameter regimes display a high density at small fractional distances, suggesting a preference for excision sites across or near the primary structure excision site as described in the maintext.

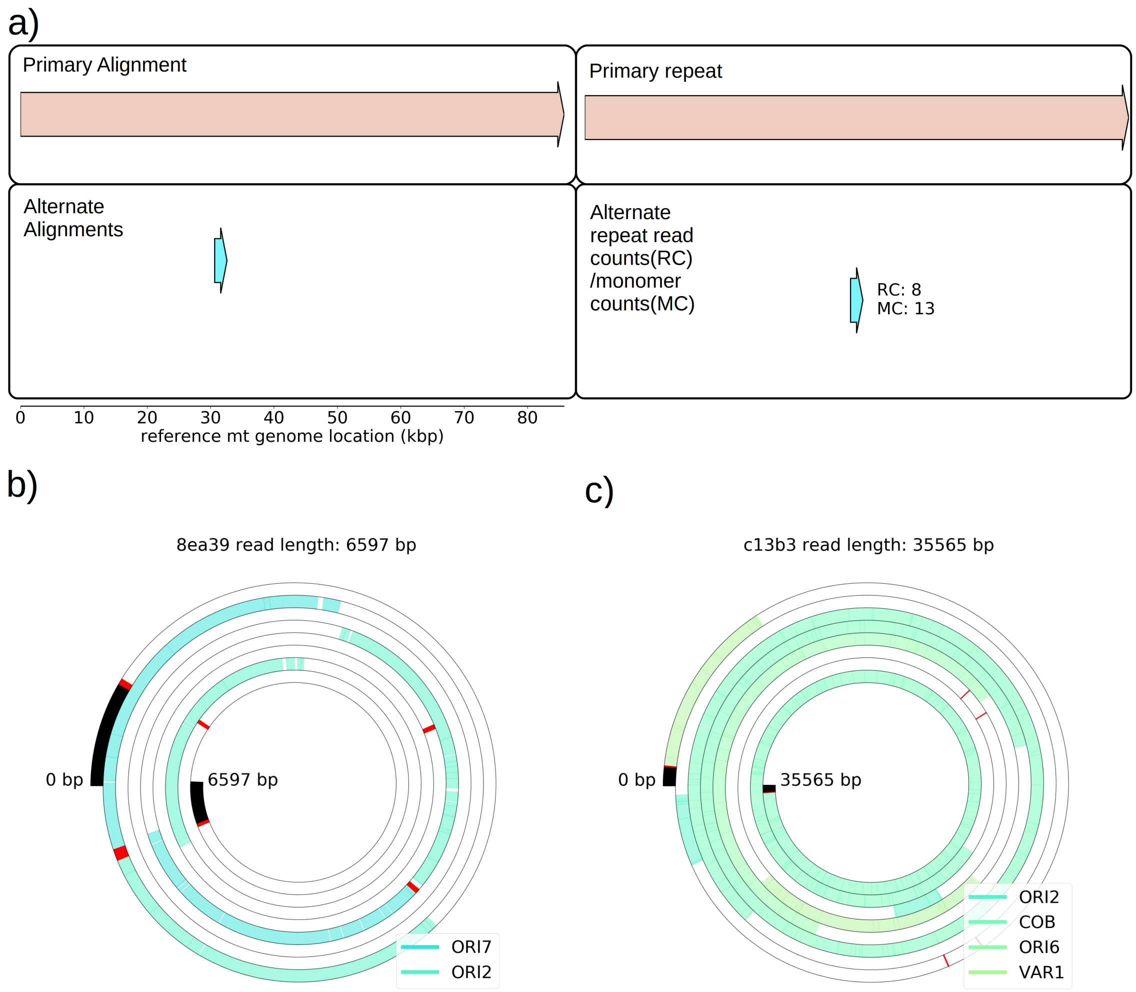


**Figure S4: Structures observed in Grande cells grown in YPG - evidence of heteroplasmy**

a) The first panel on the left shows the only high confidence primary and alternate structure detected in one of the four YPG Grande samples sequenced. Primary structure is an intact genome, while the alternate structure is a repeat that spans two origins of replication near 30kbp. The panel on the right shows the numbers of reads that support this alternate structure (RC), and the estimated number of monomer (repeat unit) counts observed across reads (MC). Given that true Petite lineages cannot exist in YPG media, this signal is due to at least transient heteroplasmy within cells. While this is the only structure detected across four samples, there are other breakpoints present at low frequencies in other YPG Grande samples, but they are not prevalent enough to infer high confidence structures from. Structure detection from Grandes is difficult because these structures are diverse (afforded by the WT genome being the reference point), and exist within cells and not lineages (therefore not enriched by chance through early bottlenecks). Given the results from (Marotta et al., 1982) that show this ori2-ori7 breakpoint is most prevalent in Petites, it is not surprising that it was only this breakpoint that was detected in YPG Grandes. b) An example read contributing to this breakpoint signal. Annotated features are coloured and labeled blocks, red lines are breakpoint locations determined from mapping. c) An example of another structure in this same YPG sample that has been detected, but is not prevalent enough for the algorithm to infer its structure (require >3 read support, and more than 2 periods of a repeat). It is a direct repeat of a region spanning from ori2 to the var1 gene.

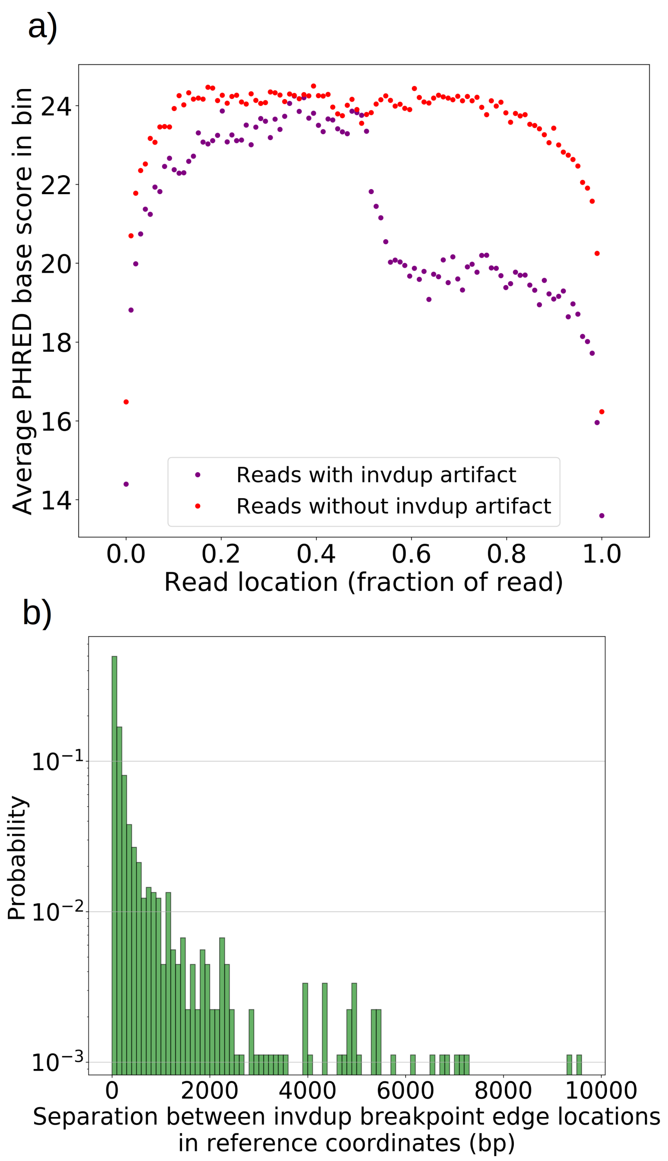


**Figure S5: Inverted duplication artifact breakpoint edges are further separated than**

**expected due to reduced basecalling accuracy**

a) For all reads with and without invdup artifacts in chrM in one representative Grande sample, a sliding window of 100bp was moved across the reads and the average PHRED scaled base quality score was computed in this window. Reads with invdup artifacts have a clear decrease in PHRED quality score in the latter half of the read due to complementary strand interaction and ratcheting as described in (Spealman et al., 2020). This decrease in base call quality means that mapping algorithms have to contend with more noise, resulting in a larger separation distance between invdup breakpoint edges which in an ideal case would perfectly coincide in reference coordinates. b) The distribution of separation distances between invdup breakpoint edges in reference coordinates. Breakpoint edges largely (90%) reside within 1 kbp of each other in reference coordinates. Inverted duplications residing outside of the expected artifact read location distribution and beyond 1 kbp separation are unlikely to be this particular sequencing artifact. We rely on these ideas in the filtering described in methods.

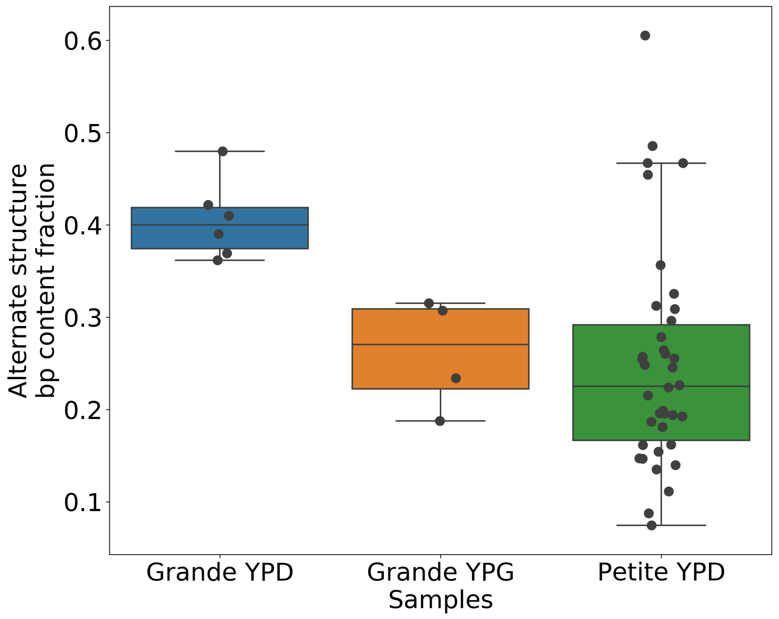


**Figure S6: Alternate structure frequencies when not neglecting reads containing inverted duplication artifacts**

In the above plot we are computing the base-pair content fraction of any reads that contain a breakpoint we have deemed to not be an inverted-duplication artifact, regardless of whether or not an artifact is present in the read. Each dot represents this alternate structure fraction across all reads in a single strain sequenced. The box plot displays the minimum, maximum, 1st quartile and 3rd quartile. We have done this same calculation for Grande samples in YPD/YPG media, and in Petites. Given that YPG samples have 2x the number of inverted duplication artifacts, which are often accompanied by spurious breakpoints, the discrepancy here between

YPD and YPG samples is strongly suggestive of clonal divergence playing a role.

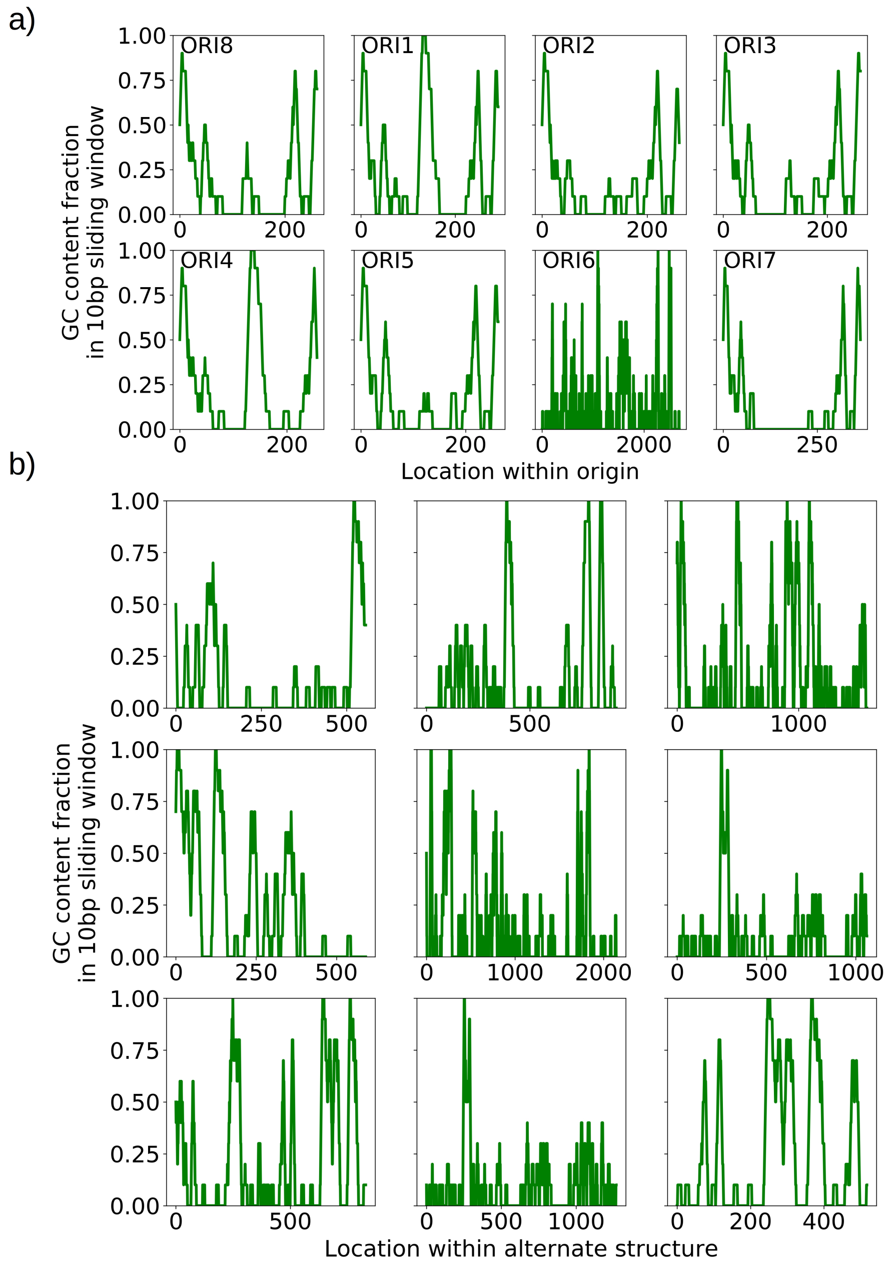


**Figure S7: GC clusters in origins and alternate structures without origins**

a) A visualization of the distinct GC clusters within all eight origins of replication. A 10bp sliding window is moved along the origin sequences. GC clusters are the 3-4 distinct peaks in each of the origin sequences, all above 0.6 GC content in the sliding window which are consistent with the locations described in (de Zamaroczy & Bernardi, 1986). b) A visualization of the GC clusters within observed alternate structures without canonical origins of replication. These clusters at similar GC content (0.6 and above) may act as surrogate replication origin sites.

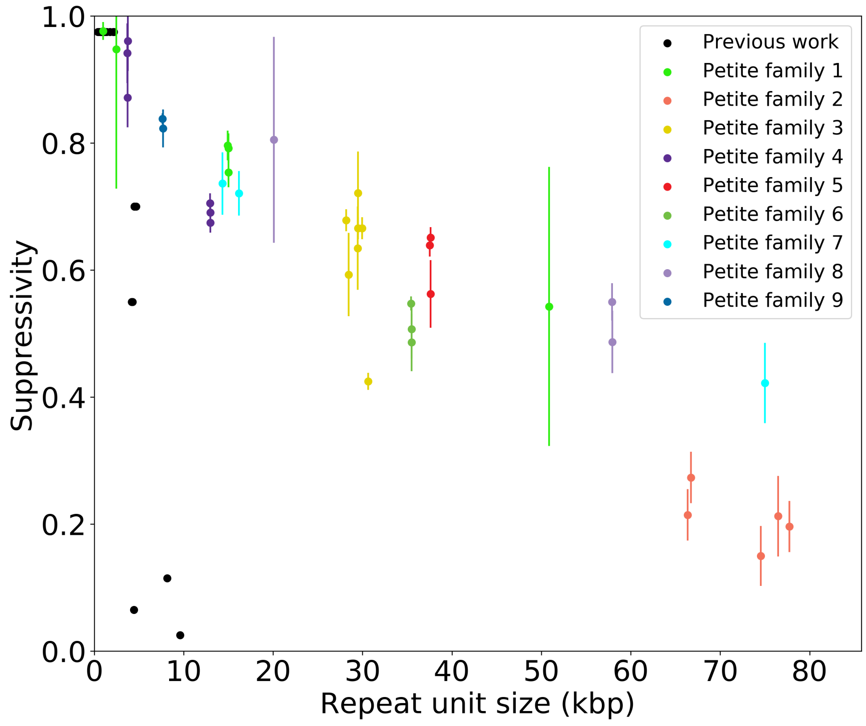


**Figure S8: Supressivity and its correlation with repeat unit size**

Black dots represent the average suppressivity and repeat unit size from the range of suppressivities and repeat unit sizes published in (de Zamaroczy et al., 1981; Mangin et al., 1983). Each coloured dot indicates the average suppressivity across three second passage Petites derived from the same first passage progenitor. Y-axis error bars are +- the standard deviation in suppresivities across these three second passage Petite colonies. Shared colours indicate that these strains were derived from the same spontaneous Petite colony and constitute a Petite family. As in the maintext, repeat unit size is taken to be the sum of the unique alignment lengths in the primary alignment of each sample. Samples containing inverted breakpoints in their primary alignments are those in family 2 and 3, the orange and yellow dots, respectively. Family 3 was confirmed to be a mixed sample.

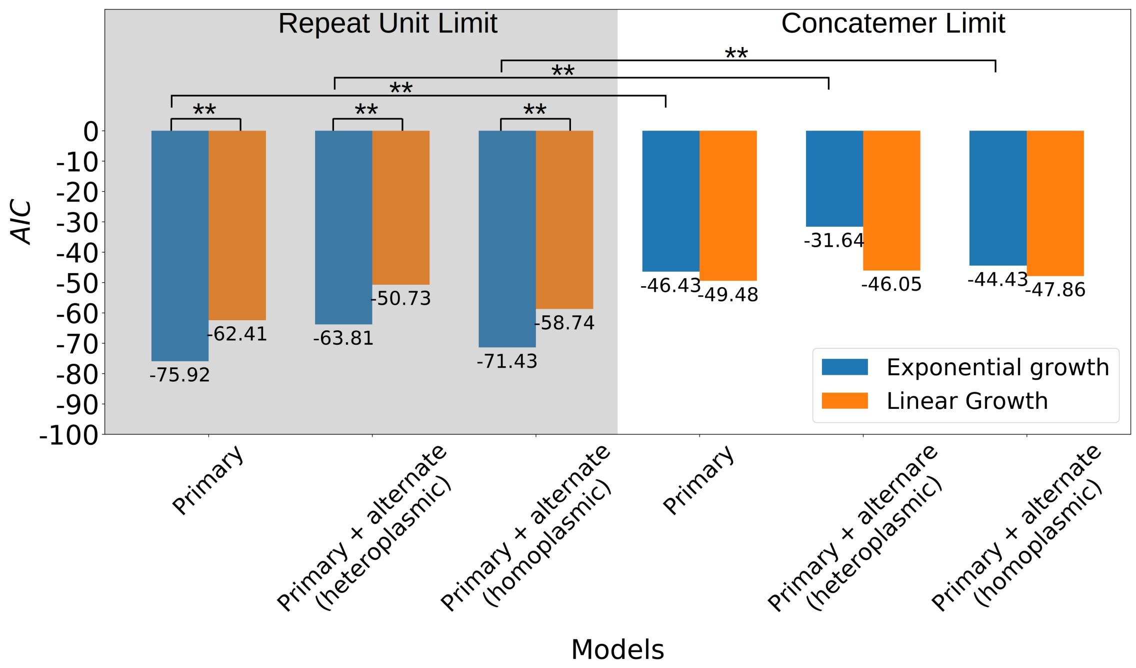


**Figure S9: Comparison of suppressivity models**

The y-axis is the Akaike Information Criterion, computed from a least-squares fit for each model by transforming the least squares statistic into a Normal negative log likelihood statistic. The left pane includes models that exist in the repeat unit limit, meaning growth rate terms ($\rho_i \nu_i$) either in the exponent (exponential), or as is (linear) are multiplied by the inverse repeat unit length ( $1/L_i$), where $i$ is Petite or Grande. These products represent the time evolution of the abundance of each structure ( $N_i$ as shown in the axis of Figure 8 in the maintext). The right pane is the concatemer limit, which takes the same form of the models in the left except with the exclusion of the inverse repeat unit length prefactors. The notation ‘Primary’ indicates that only primary structures are considered in the theoretical suppressivity calculation, while ‘Primary + alternate’ indicates that both alternate structures and primary structures observed in strains contribute to the theoretical suppressivity. There are two ways alternate structures are included in the models here: (I) The heteroplasmic limit, where it is assumed these structures coexist. In this case, the relative contributions of alternate/primary structures are included by computing $N_P$ as an average over all structures weighted by their mitochondrial contributions. (II) The homoplasmic limit, where it is assumed all structures are segregated into their own homoplasmic lineages. In this case, the theoretical suppressivities are an average weighted by mitochondrial contributions of each structure. By comparing the relative likelihood of each model, in the repeat unit limit the exponential model is significantly favoured over the linear model (**, 2$\sigma$). All pairwise comparisons between the same models in the repeat unit limit are significantly favoured over the concatemer models with the exception of the heteroplasmic linear models. Homoplasmic/heteroplasmic limits with the inclusion of alternate structures has little effect.

a)

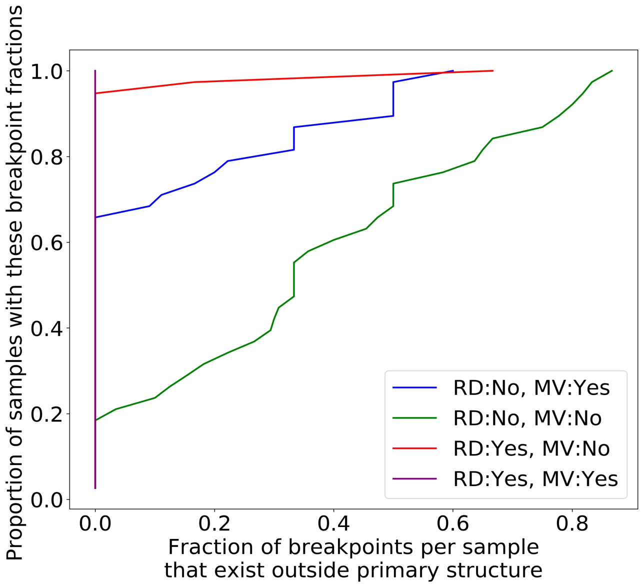


b)

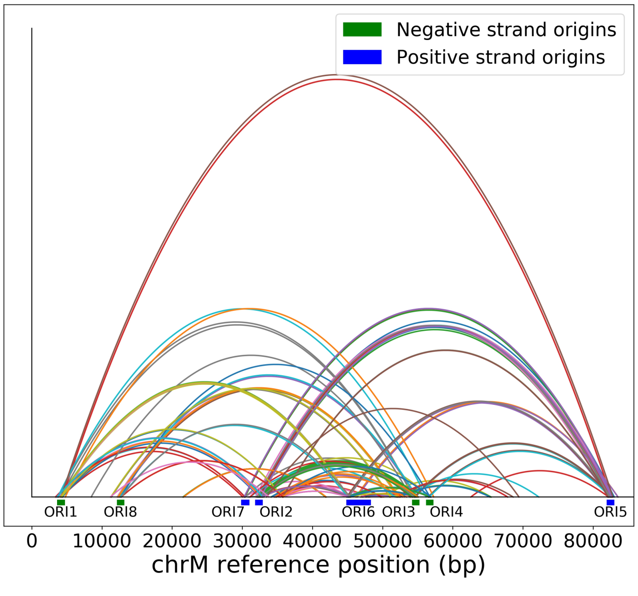


**Figure S10: Effect of repeat detection (RD) and majority voting scheme (MV) on breakpoint filtering**

a) Between the green and blue curve we see the effect that majority voting (MV in legend) has on the cumulative fraction of breakpoints that exist outside the primary alignment (and are therefore unlikely to be real given the notion of an excision cascade). Between the green and red curve we see the effect of tandem repeat detection (RD in legend) of breakpoint labels in reads, which largely eliminates breakpoints that exist outside of the primary alignment. The purple curve indicates the effect of both repeat detection and majority voting, indicating that across all samples breakpoints are contained within the primary alignment which makes them believable. b) Here we are plotting the locations of breakpoints that are removed through repeat detection and the majority voting process. Breakpoints that are removed this way largely bridge between origins of replication due to significant homology. Sequencing errors can perturb one origin into another, resulting in these spurious breakpoints that are removed by these two schemes.

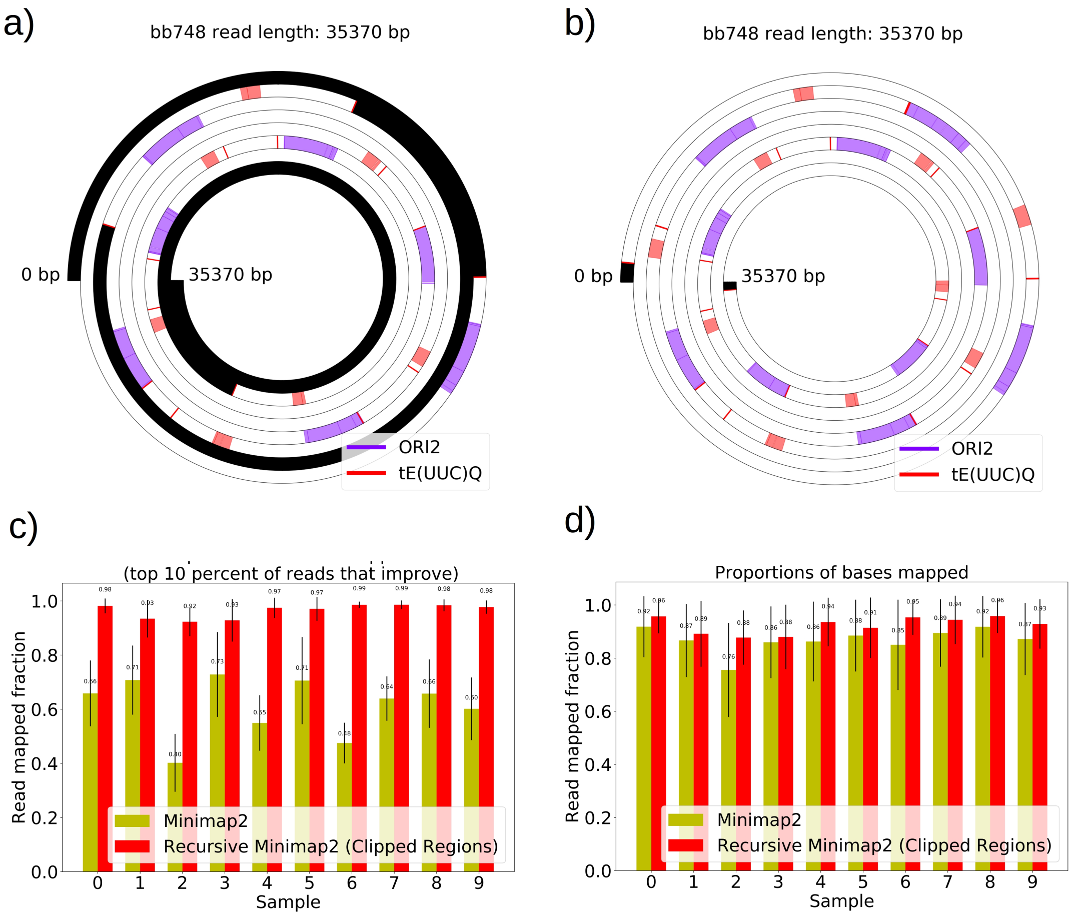


**Figure S11: The necessity and effect of recursive Minimap2 alignments in resolving Petite structure**

a) A spiral plot showing a raw Minimap2 alignment of a Nanopore read with default parameters. The legend indicates annotated features that are present, black regions represent clipped (unmapped) regions, red bars indicate the start or ends of adjacent alignments. Large portions of this read are unmapped due to the default Minimap2 z-offset parameter, which truncates repeated alignments due to the expectation of colinearity with the reference sequence. Instead of varying this parameter, which requires balancing early truncation and enforcing colinearity with the reference, we opted to recursively apply Minimap2 in unmapped portions after the first run. b) The effect of recursively mapping unmapped portions in the same read which almost entirely eliminates the unmapped regions except at the ends of reads where adapters still reside and sequencing error is generally higher. While this produces a pseudo-global alignment, only alignments with MAPQ > 20 are retained in subsequent analysis so our requirements of alignment specificity are maintained. c) The most impactful change that recursive mapping has in improving mapping fraction in 10 samples. d) The global effect, which is minimal, but does improve statistics in repeat detection and structure construction especially in low frequency structures where every read counts.

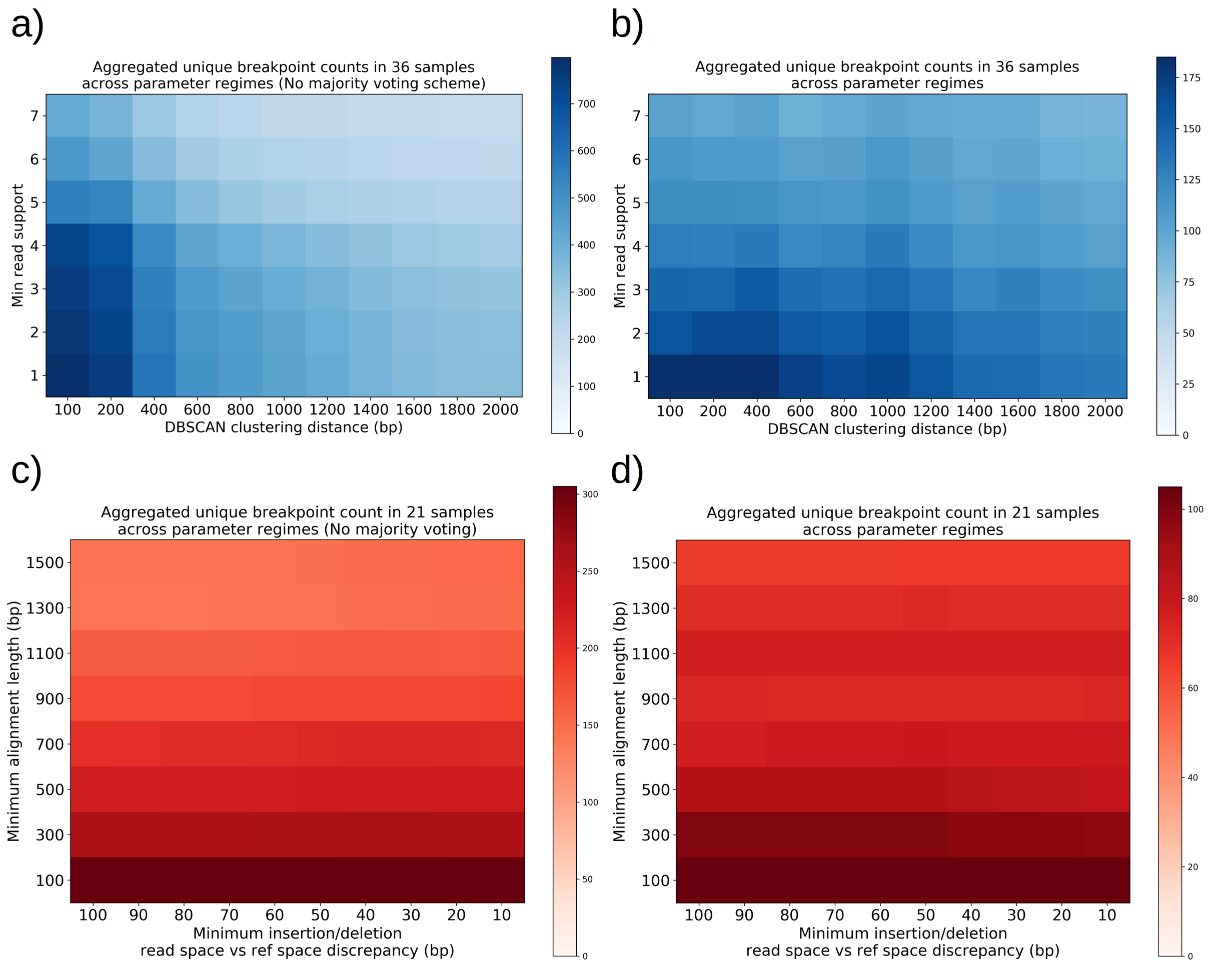


**Figure S12: Structural detection pipeline parameter sweeps - The effect of majority voting, minimum read support, and DBSCAN clustering radius, insertion/deletion size threshold, and minimum alignment length**

a) A plot of total unique breakpoint counts (identified through DBSCAN clustering) without applying the majority voting scheme described in methods across a parameter sweep of DBSCAN clustering radius and minimum read support. Minimum alignment length and minimum insertion/deletion size were fixed at 300bp, and 30bp, respectively. As expected, we see a monotonic decrease in counts as clustering radius is expanded, and the same type of decrease for increasing read support. The high density cluster to the lower left is an indication that in this regime we are largely clustering noise. b) The same plot in panel (a) but with majority voting for breakpoints, which results in a 4 fold decrease in unique breakpoint counts at the extremum. The flatness across these parameter ranges is a sign that the breakpoint counts represent real structures that are largely insensitive to parameter selection with the exception of a small clustering radius which will always produce more clusters in the lower left. c) A plot of total unique breakpoint counts (identified through DBSCAN clustering) without applying the majority voting scheme described in methods across a parameter sweep of minimum insertion/deletion length and minimum alignment length. The DBSCAN clustering radius and minimum read support were fixed at 1kbp, and 3, respectively. We see low variance in counts along the insertion/deletion length threshold axis, suggesting that most breakpoint calls from minimap2 on its own are capturing real breakpoints and that this parameter has little effect. Increasing minimum alignment length as expected results in a monotonic decrease in breakpoint counts because less reads exist in the tail of the read-length distribution, and because small structures are thrown away. d) The same plot in panel (c) but with majority voting for breakpoints, which results in a 3-fold decrease in unique breakpoint counts at the extremum. While we see less steep descent along the minimum alignment length axis, it is clear that below 300bp in alignment length there appears to be a transition to clustering of noise, which is due to spurious replication origin to replication origin transitions (Figure S10).

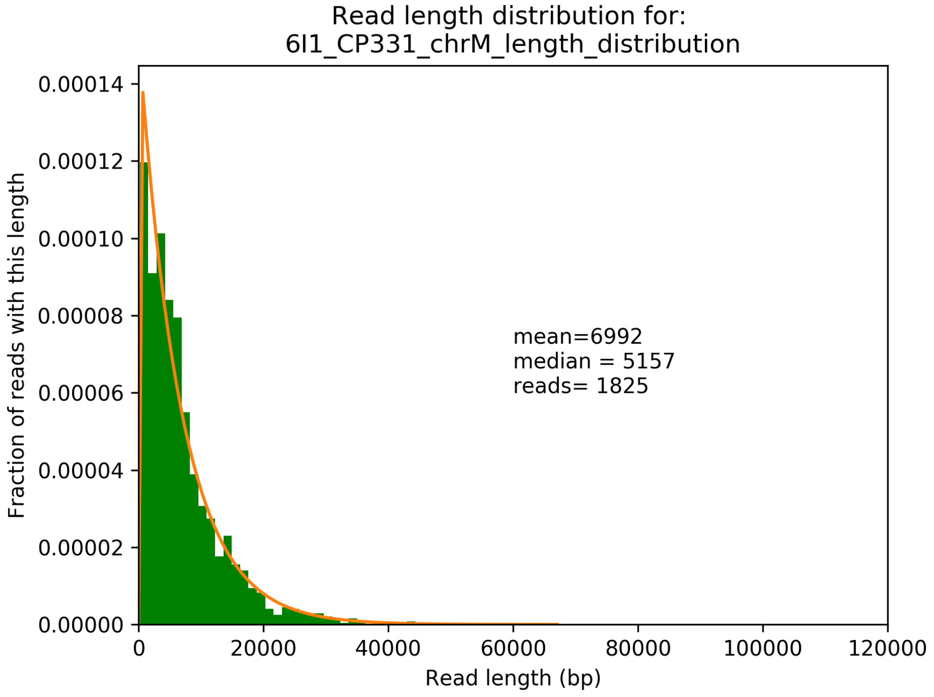


**Figure S13: Example read length distribution from sequencing of a single Petite strain and its exponential fit**

A plot of an empirical read length distribution from one sequence Petite strain. Mean, median, and the number of reads are denoted in addition to a fit to an exponential probability distribution in orange. For this particular fit, the location and scale parameters were 213, and 6779, respectively.

### Supplementary Information: Tables

#### Table S1: Structural detection pipeline overview and summary of parameters

*Summary of algorithm*

1.) Dataset across all samples is restricted to reads that pass default Guppy QC, alignments with PHRED alignment score >20 and alignment length > 300bp.
2.) Breakpoints in alignments where the read-space and reference-space differ by >30bp are recorded as “breakpoints”.
3.) Inverted breakpoints, where the strand changes mid read, are filtered from hairpin artifacts based on the positions within reads, allowing for 1% false positive error at this stage (see Figure S1). Hairpin artifacts are marked for use later in a majority voting scheme.
4.) Non-inverted breakpoints, and inverted breakpoints are separately clustered using DBSCAN, with a clustering distance of (eps) of 1kbp, and a minimum cluster occupancy of 3.
5.) All breakpoints are further filtered with a majority voting scheme which eliminates clusters that exist due to sequencing error and are not periodic (assuming concatemer structures), or are not recapitulated on either strand of read when both strands are present. The latter is reminiscent of Oxford Nanopore 2D basecalling, but is performed post alignment.
6.) Breakpoints are further filtered so that only those represented across > 3 separate reads remain.

*Detailed summary of parameters*

| Parameter | Value | Description/justification of choice of value |
|---|---|---|
| PHRED alignment score | 20 | 1/100 probability of a false positive alignment by chance given a particular size/complexity of reference |
| Minimum alignment length | 300bp | Mt replication origins are ~300bp on average and exhibit significant homology. Below 300bp, alignments containing origin fragments are often indistinguishable in Nanopore sequencing error background, resulting in erroneous alignments that break expected |

| | | |
|---|---|---|
| | | collinearity. This would also be the expected lower limit for detectable repeat units in Petites. See Figure S12 for effects of this parameter. |
| Insertion/deletion threshold | 30bp | The minimum discrepancy in reference vs read space to call a breakpoint. This parameter appears to have little effect at low values, suggesting breakpoints we see break collinearity by much larger distances, and are therefore more believable over Nanopore sequencing error background which often introduces small (~10bp) insertions and deletions. See Figure S12 for effects of this parameter. |
| DBSCAN epsilon (minimum clustering distance) | 1kbp | This is the upper threshold for Sniffles and NanoSV clustering, and will cluster within smaller distances if the SV's themselves are smaller. Varying this value has little effect even in a range of a few hundred bp due to the majority voting scheme. See Figure S12 for the effect of this parameter. |
| DBSCAN minimum cluster occupancy | 3 | Fixed at 3 in this pipeline to allow for small clusters, and generally affected by coverage. More stringent filtering along similar lines is available with read support filtering. Also important to note that we don't see high confidence structures with breakpoint counts < 10 once repeats have been detected. |
| Minimum read support | 3 | Reasonable values for this parameter are largely dependent on sequencing coverage. See Figure S12. |

**Table S2: Definitions of parameter regimes referenced in Figure S2 and Figure S3**

| **Parameter** | **"Lenient" parameter set value** | **"Selected" parameter set value** | **"Strict" parameter set value** |
|---|---|---|---|
| Minimum alignment length | 100bp | 300bp | 1kbp |
| Insertion/ | 30bp | 30bp | 30bp |

| | | | |
|---|---|---|---|
| deletion threshold | | | |
| DBSCAN epsilon (minimum clustering distance) | 100bp | 1kbp | 1.4kbp |
| Minimum read support | 1 | 3 | 5 |
| Majority voting? | no | yes | yes |
